# Lifestyle, Early-life, and Genetic Health Risk Factors Underlying the Brain Age Gap: A Mega-Analysis Across 3,934 Individuals from the ENIGMA MDD Consortium

**DOI:** 10.1101/2025.05.09.653064

**Authors:** Laura K.M. Han, Yara J. Toenders, Xueyi Shen, Yuri Milaneschi, Heather C. Whalley, Philipp G. Sämann, Til Andlauer, Jochen Bauer, Klaus Berger, Tiana Borgers, James H. Cole, Udo Dannlowski, Kira Flinkenflügel, Hans J. Grabe, Oliver Gruber, Tim Hahn, Paul J. Hamilton, Sean N. Hatton, Marco Hermesdorf, Ian Hickie, Jan Homann, Tilo J. Kircher, Bernd Krämer, Anna Kraus, Axel Krug, Christina M. Lill, Sarah E. Medland, Susanne Meinert, Alana Panzenhagen, Brenda W.J.H. Penninx, Nic J.A. van der Wee, Marie-José van Tol, Uwe Völker, Henry Völzke, Antoine Weihs, Katharina Wittfeld, Sophia I. Thomopoulos, Neda Jahanshad, Paul M. Thompson, Elena Pozzi, Lianne Schmaal

## Abstract

**Background:** Large-scale studies show that adults with major depressive disorder (MDD) generally have a higher imaging-predicted age relative to their chronological age (i.e., positive brain age gap) compared to controls, though considerable within-group variation exists. This study examines lifestyle, early-life, and genetic health risk factors contributing to the brain age gap. Identifying risk and resilience factors could help protect brain and mental health.

**Methods:** Using an established model trained on FreeSurfer-derived brain regions (www.photon-ai.com/enigma_brainage), we generated brain age predictions for 1,846 controls and 2,088 individuals with MDD (aged 18-75) from 12 international cohorts. Polygenic risk scores (PRS) were calculated for major depression, C-reactive protein, and body mass index (BMI) using large-scale GWAS results. Linear mixed models were applied to assess lifestyle (BMI, smoking, education), early-life childhood trauma, and genetic (PRS) health risk associations with the brain age gap. Additionally, we evaluated the link between the brain age gap and peripheral biological age indicators (epigenetic clocks).

**Results:** Higher brain age gaps were significantly associated with BMI (β=0.01, P_FDR_=0.02) and smoking (β=0.11, P_FDR_=0.02), while lower brain age gaps were linked to higher education (β=-0.02, P_FDR_=0.02). Higher childhood trauma scores predicted a higher brain age gap (β=0.04, P=0.01). Higher brain age gaps were positively associated with all PRS (βs=0.04-0.16, Ps_FDR_=0.02-0.03). There were no significant interactions between diagnosis and assessed factors on the brain age gap. In a multivariable model, only modifiable health factors—BMI, smoking, and education—remained uniquely associated with brain age gaps.

**Conclusions:** Genetic liability for depression and related traits is linked to poorer brain health, but health behaviors potentially offer a key opportunity for intervention. This study underscores the importance of targeting modifiable lifestyle factors to mitigate poor brain health in depressed individuals, an approach perhaps under-recognized in clinical practice.

## INTRODUCTION

Individuals with major depressive disorder (MDD) are often more vulnerable to age-related changes and experience poorer physical health than the general population.^1–5^ As a result, the global burden of MDD transcends the false dichotomy between mental and physical health.^6^ Previous research has identified complex genetic-brain-behavioral disturbances in MDD,^7^ including those linked to age-related biological signatures.^8,9^ A large body of evidence demonstrates the multifactorial nature of MDD, highlighting the interplay of genetic, environmental, psychological, and biological dysregulations that contribute to its symptomatology.^10^ This complex interplay helps explain why depression is often accompanied by a wide range of physical comorbidities and somatic consequences.^11,12^ By improving the prevention, early detection, management, and treatment of MDD, we have the potential to achieve substantial gains in overall health and reduce healthcare costs.^13^

Accumulating evidence suggests that, at the group level, individuals with MDD follow advanced aging trajectories.^14–18^ Their functional (e.g., walking speed, handgrip strength) and biological state (e.g., telomeres, mitochondrial function) often reflects what is typically seen in older individuals, suggesting that their biological age may “outpace” their chronological age, though much of this evidence comes from cross-sectional studies. The “brain age” paradigm has gained significant traction over the past decade as a method to assess the biological age of the brain.^19,20^ This approach posits that a larger gap between chronological age and predicted brain age is associated with unfavorable clinical outcomes in patients with neuropsychiatric disorders.^21^ Using this framework, our research, along with others, has shown that individuals with MDD indeed exhibit a higher brain age gap compared to non-depressed peers.^22–27^ However, it is important to mention the large within-group and small between-group variance, demonstrating that many patients do not show abnormal age-related structural brain patterns. This raises the question of *when* patients begin to deviate from the typical brain age trajectory and *what* factors contribute to the individual variations in the brain age gap between depressed and healthy populations.

In the present study, we leverage the large-scale ENIGMA Consortium MDD Working Group^28^ to bring together multiple methodologies to connect individual brain signatures with genetic risk profiles and health risk factors, aiming to investigate their unique and shared contributions to brain health. Specifically, we focus on age-related structural brain patterns, assessing the relationship between the brain age gap and several factors that, while measured cross-sectionally, may indirectly reflect aspects of an individual’s life history. First, we will examine lifestyle factors—such as body mass index (BMI), smoking behavior, and education. BMI and smoking have been commonly reported to be associated with age-related structural brain differences, as well as potential protective effects of educational attainment.^29–34^ Next, we investigate childhood trauma as an early-life environmental influence. Previous studies, albeit with different imaging modalities^35^ or in young developing individuals (<18 years), suggest that different types of early-life adversities are differentially associated with the brain age gap.^36,37^ Finally, we will assess polygenic risk scores (PRS) as proxies for inherited health risk predispositions. A PRS is an individualized estimate of genetic liability to a trait or disease, calculated as the sum of genetic susceptibility variants weighted by their effect sizes derived from independent genome-wide association study (GWAS) results.^38^ While one study identified a weak but common genetic architecture between regionally estimated brain age gaps and MDD,^39^ GWAS conducted in UK Biobank found no genetic correlation or mendelian randomization causal estimates between the brain age gap and MDD.^40,41^ A key question we address here is whether individual variation in the brain age gap can be explained by differences in (genetic liability for) MDD and MDD-related traits, or whether it is primarily associated with a combination of indirect or confounding factors.

Despite growing interest in the brain age paradigm, methodological heterogeneity across studies has led to inconsistent findings regarding the factors that contribute to the brain age gap.^42^ To address this, we deploy a previously validated brain age prediction model across 3,934 individuals from 12 cohorts, to determine robust associations between the brain age gap and a variety of genetic and health risk and resilience factors. We also examine whether these relationships are differentially affected by diagnostic status, given the substantial overlap in brain age gap variability observed in both controls and individuals with MDD. Based on this overlap, we hypothesized that the associations between the brain age gap and the PRS and health risks would not differ between cases and controls. Additionally, we sought to identify which health risk factors are most strongly and uniquely associated with the brain age gap, using a multivariable model. Finally, to further assess the relationship between peripheral and brain imaging age-related signatures, we conducted exploratory analyses to examine associations between the brain age gap and (PRS for) several DNA methylation (DNAm)-derived biological age indicators (epigenetic clocks).^43–46^ Together, these analyses aim to provide a more comprehensive understanding of the factors involved in individual brain age variations and their implications for MDD.

## METHODS

### Participants

A total of N=3,934 individuals (N=1,846 controls and N=2,088 persons with MDD) aged 18-75 years old with European ancestry from 12 cohorts of the ENIGMA MDD working group participated in this study. MDD was ascertained using diagnostic interviews. Cohort-specific details on demographics, basic clinical characteristics, and exclusion criteria can be found in **Supplementary Table S1**. Individual cohorts obtained approval from their local institutional review boards and ethics committees. Written informed consent was provided by all study participants.

### Image processing and analysis

Structural 3D volumetric T1-weighted brain MR images of each subject were acquired at each site. Standardized protocols were used to facilitate harmonized image analysis and feature extraction (N=153) across multiple sites (http://enigma.ini.usc.edu/protocols/imaging-protocols/). Within FreeSurfer,^47^ the Desikan/Killiany atlas was used for the cortical parcellations and the *aseg* atlas was used to segment 14 subcortical gray matter regions. The nucleus accumbens, amygdala, caudate, hippocampus, pallidum, putamen, thalamus, two lateral ventricles, 68 cortical thickness and 68 cortical surface area measures, and total intracranial volume (ICV) were statistically examined for outliers and visually inspected. The **Supplementary Table S2** includes more detailed information on the image acquisition parameters, software descriptions, and quality control.

### Brain age prediction model

The model development is detailed in Han et al. (2021, 2022)^22,24^ but briefly, ridge regression was used to predict age from 77 FreeSurfer-derived features in healthy controls without a history of mental or neurological illness. FreeSurfer features were averaged across hemispheres. Separate models were trained for male (N=952) and female (N=1,236) controls. The model used to generate brain age predictions is publicly available (www.photon-ai.com/enigma_brainage) and was applied to the ENIGMA MDD cohorts included in the current study. The brain age predictions in the current dataset have previously been generated, evaluated, and published.^22,24^ Model performance (mean absolute error, weighted MAE, Pearson correlation coefficients between predicted brain age and chronological age, and proportion of the age variance explained by the model *R^2^*) in the current dataset can be found in the **Supplementary Table S3**. Participants used to train the brain age prediction models were excluded from the current study. While the performance of the current model may be suboptimal compared to more recently developed brain age prediction models that, for example, use raw MR images, a significant advantage of the current approach is its use of FreeSurfer-derived features as input. This enables anonymized data sharing, facilitating much larger sample sizes than studies using models that rely on raw images. Large sample sizes are crucial for reliably estimating brain-genetics-behavior associations.

### GWAS summary statistics

For this study, we used the summary statistics from three large discovery GWAS to calculate PRS for major depression including established MDD diagnosis or self-declared depression,^48^ BMI,^49^ and C-reactive protein (CRP).^50^ Despite the broader definition of depression, the GWAS used here demonstrates high genetic correlations with a clinically ascertained MDD GWAS (rG = 0.86).^48^ The MDD GWAS summary statistics used in the current study did not include data from 23andMe. For all three GWAS summary statistics, all overlapping individuals with the ENIGMA MDD datasets were excluded. To identify and remove any potential overlapping individuals between the discovery GWAS of MDD and testing samples, we used the Psychiatric Genetic Consortium (PGC) CheckSums algorithm with the following approach.^48,51^ First, the CheckSums scripts (https://personal.broadinstitute.org/sripke/share_links/checksums_download/) were shared with participating ENIGMA MDD cohorts. Second, analysts at the participating site ran these scripts on their data and returned a key to uniquely identify genotypically identical individuals without sharing any genotype configuration. Finally, the output was subsequently shared with the PGC GWAS analyst. Following this approach, no participating cohort within ENIGMA MDD had access to the CheckSums data of PGC nor any other ENIGMA MDD cohort. Any participant that was both part of the GWAS and the ENIGMA MDD consortium was removed from the discovery GWAS and an updated analysis of the summary statistics was carried out by the PGC analyst to ensure there was no overlap between the discovery and test samples. For the BMI GWAS summary statistics, we removed all overlapping cohorts. The final MDD GWAS summary statistics covered 727,742 individuals. The final BMI GWAS summary statistics covered 331,616 individuals. The final CRP summary statistics covered 204,402 individuals.

### Genetic data

Preprocessing and quality checking were conducted locally by each participating cohort. Only the anonymized individual-level PRS data were shared for analyses. Imputed, hard-call, genetic data were used to generate PRS. To select the SNPs for creating PRS, a two-stage approach was applied. First, all ENIGMA cohorts provided a list of SNPs from the hg19/GRCh37 build that passed QC criteria of a minor allele frequency (MAF) > 0.01 and INFO scores > 0.1. Subsequently, three SNP lists were created for each individual cohort to generate PRS: 1) a hard list: SNPs present in all ENIGMA MDD cohorts (N_SNP_ = 3,176,977), 2) a soft list: SNPs present in >80% of the ENIGMA MDD cohorts (N_SNP_ = 6,306,997), and 3) a cohort list: all SNPs that passed QC for that specific cohort (N_SNP_ varies per cohort).

### Calculation of PRS

To calculate PRS, we used a clumping and thresholding method (PRS-CT).^52^ For each phenotype (i.e., MDD, BMI, and CRP) ten PRS were calculated using PRSice 2.0 based on ten p-value thresholds (pT = 5×10^-8^, 1×10^-6^, 1×10^-4^, 1×10^-3^, 0.01, 0.05, 0.1, 0.2, 0.5 and 1). A window of 500kb base pairs with an r^2^ <0.1 was used to conduct the clumping. Genotype data of central European samples (CEU) from the 1000 Genome Project were used as a reference panel for clumping. The scripts used to calculate the PRS are publicly available at https://github.com/xshen796/ENIGMA_mdd_prs/blob/main/script/PREP_PRS/Calculate_PRS.md.

### Validation of PRS prediction

To validate the PRS, out-of-sample prediction of the PRS at all thresholds (ten PRS per phenotype/trait in total) was performed using linear mixed models. Covariates that account for genetic relatedness and population stratification were residualized from the PRS by regressing them against genomic relationship matrices (GRM). The GRM were calculated by using the imputed genetic data that passed the quality check using GCTA.^53^ The first ten genetic principal components (PCs) were included as additional covariates. Mixed-effect logistic regression was used to examine the association between the residualized PRS and MDD diagnosis using the ‘glmer’ function in the ‘lme4’ R package, by taking MDD diagnosis as the dichotomous outcome and the PRS for MDD as the predictor. Similarly, a linear mixed model using the ‘lme’ function in the ‘nlme’ R package was used to validate the PRS for BMI, by taking BMI as a continuous outcome and the PRS for BMI as predictor. In both models, age and sex were included as fixed effects, and scanning site as a random factor. The PRS for CRP has been previously validated,^50^ but no phenotypic CRP data was available to perform validation analyses in the current study.

### Health risk and resilience factors

Childhood trauma was measured using the total Childhood Trauma Questionnaire short form (CTQ-SF) score.^54^ The CTQ-SF is a reliable and valid self-report instrument ^55^ consisting of 28 items divided over five subscales of childhood adversity: emotional, physical, and sexual abuse, and emotional and physical neglect that sum up to a total score. The CTQ total score was used as a continuous variable. BMI (kg/m^2^) and current smoking status (yes/no) were assessed at time of scanning. Education was assessed as the total number of years of completed education (school/university/vocational training) and also used as a continuous variable.

### Statistical analyses

Individual-level brain age gap scores, PRS, and demographic and health risk measures from each cohort were pooled for a mega-analysis. PRS scores were adjusted for population stratification using the first ten genetic PCs, and brain age gap scores were adjusted for both linear and quadratic age effects in all models. Sex was included as a covariate throughout. To account for site-related variability, random intercepts for scanning sites (N=17) were incorporated into all models. First, we used linear mixed models to examine the associations between the brain age gap and the following factors:

1. Lifestyle factors (BMI, education, and current smoking status)
2. Early-life factor (Childhood trauma)
3. Genetic factors (PRS for MDD, BMI, and CRP)

These associations were analyzed across individuals with and without major depressive disorder MDD. Second, we tested interaction terms between the predictor of interest and MDD status (yes/no) to determine whether MDD moderated the relationship between the brain age gap and the respective lifestyle, early-life or genetic risk factor. Third, significant health risk factors were included in one multivariable model to examine which factors were most strongly and uniquely associated with the brain age gap. Sensitivity analyses were performed by adding FreeSurfer version as an additional covariate in the models to examine whether findings were independent of the preprocessing pipeline used.^24^ Continuous variables with values in the lowest or highest 5% were winsorized to the respective 5th and 95th percentile thresholds. All continuous variables were scaled and centered prior to inclusion in the models. Standardized regression coefficients were reported as effect sizes. All tests were two-sided, and P-values were adjusted for multiple comparisons within each corresponding domain (lifestyle: across 3 factors and genetic: across 3 PRS) using the false discovery rate (FDR) method. Findings were deemed statistically significant at an FDR-adjusted P-value threshold of <0.05.

### Exploratory analyses

To investigate the relationship between peripheral and central indicators of biological age, we also explored associations between the brain age gap and PRS for several epigenetic clocks (i.e., GrimAge, Hannum age, Horvath age, and PhenoAge acceleration). While Hannum (based on blood)^44^ and Horvath (based on multiple somatic tissues)^43^ epigenetic clocks are trained to predict age from DNAm data (“first-generation”), GrimAge^45^ and PhenoAge^46^ epigenetic clocks (both based on blood) are optimized to enhance the prediction for aging- and mortality-related outcomes (“second-generation”). The GWAS summary statistics were derived from more than 40,000 individuals and are based on age-adjusted residuals of the epigenetic clocks (i.e., age acceleration).^56^ DNAm data were available for two cohorts (N=810; chronological age ranging from 20 to 75 years), allowing us to: a) validate the PRS predictions, and b) examine the relationship between the brain age gap and epigenetic clock phenotypes within these cohorts. The analyses in b) were corrected for linear and nonlinear age effects, sex, and the time difference between blood draw and MRI scan which varied for one cohort. Random intercepts were used to account for scanner differences. Batch effects were regressed from the epigenetic clock phenotypes prior to inclusion in the model. Finally, we tested the associations between the brain age gap and PRS for the epigenetic clocks across all cohorts with genetic data.

## RESULTS

### Participant characteristics

The demographics and assessed phenotypes of the study sample are summarized in **Table 1**. Briefly, the MDD group, on average, had an older chronological age and a higher proportion of females compared to the control group. As expected, the MDD group also had a higher average BMI, higher proportion of current smokers, increased exposure to childhood trauma, and fewer years of education. As previously demonstrated using these samples,^22,24^ we confirm that in this subset of individuals with both imaging and genetic data, those with MDD exhibited a higher brain age gap compared to controls. This difference was observed both in raw comparisons and after adjusting for age, age², sex, and random intercepts for site (b = +1.06 years [SE=0.22], p<0.0001; CI 95% 0.07-0.19; Cohen’s d = 0.13 [SE=0.03]).

**Table 1.**
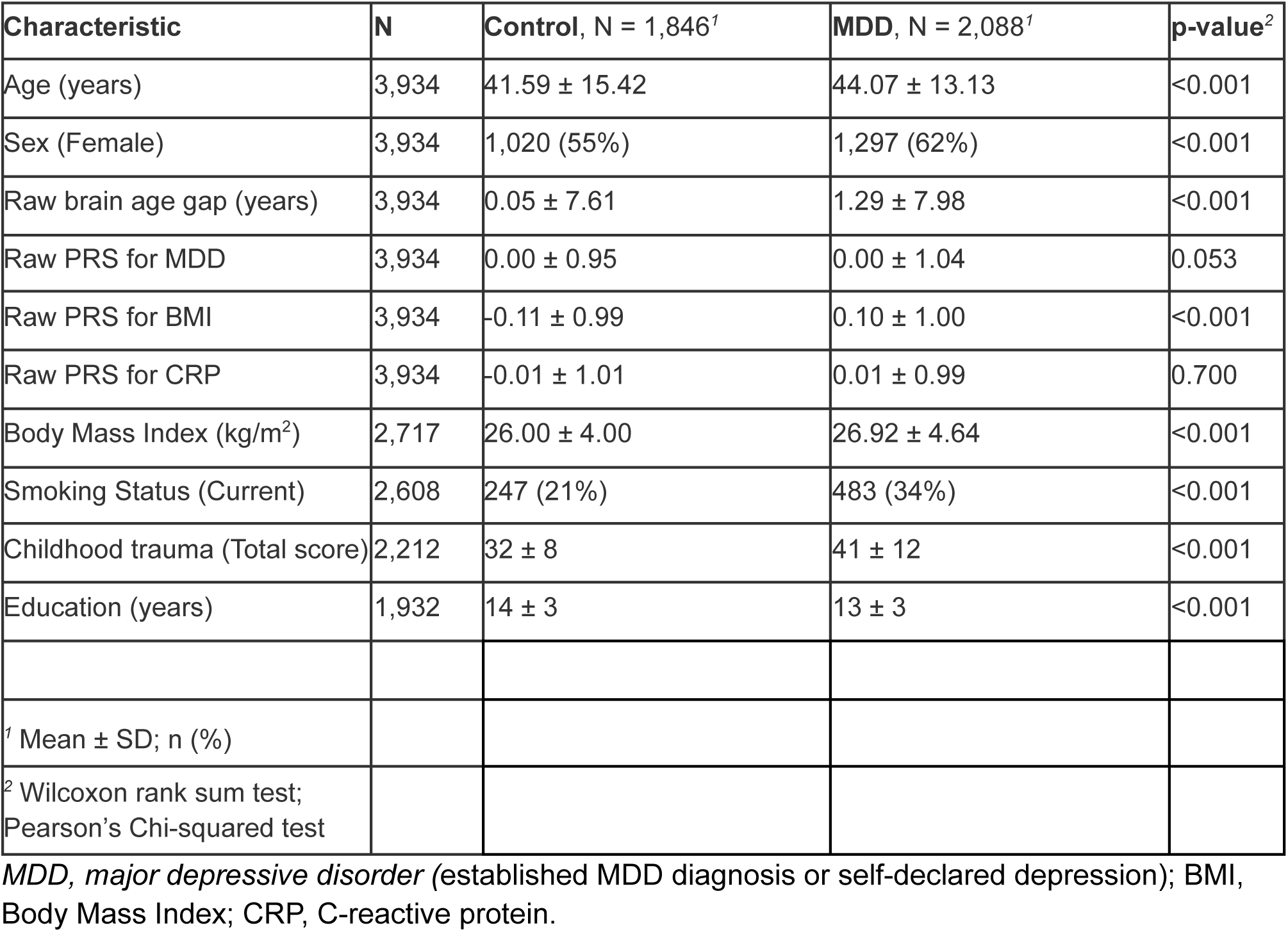
Participant Characteristics.

### Validation of PRS prediction

The PRS calculated using the three different SNP lists were highly correlated (all r > 0.99). The analyses were therefore conducted using the “cohort list” variant that was based on the SNPs that passed QC criteria for each individual cohort. For the validation of the PRS for MDD, all PRS created using p-value thresholds equal to or higher than 0.0002 were significantly associated with MDD diagnosis with odds ratios ranging from 1.04 to 1.17 and P-values ranging from 6.4×10^-5^ to 0.039 (**Supplementary Table S4**). For the validation of the PRS for BMI, all PRS created at all ten thresholds were significantly associated with BMI with β-values ranging from 0.07 to 0.19 and P-values ranging from 8.0×10^-26^ to 0.0002 (**Supplementary Table S5**). For the PRS for CRP, no phenotypic CRP data was available to perform validation analyses.

### Association between the brain age gap and lifestyle and early-life factors

A multivariable model indicated that all considered lifestyle factors were significantly associated with the brain age gap (all **Ps_FDR_** < 0.02). First, each 1 standard deviation (SD) increase in BMI is associated with a 0.01 SD increase in the brain age gap, and current smokers exhibited, on average, a 0.11 SD higher brain age gap than non-smokers. In terms of resilience factors, each 1 SD increase in education was associated with a -0.02 SD decrease in the brain age gap. Second, each 1 SD increase on the total score of the CTQ was associated with a 0.04 SD higher brain age gap. An overview of these findings is provided in **Table 2**. Sensitivity analyses, which included the FreeSurfer version used for preprocessing as an additional covariate in the model, did not alter any of the findings.

**Table 2.** Associations between the brain age gap and risk and resilience factors.

| Predictor | $\beta$ | SE | lower CI | upper CI | DF | t-value | P-value | P<sub>FDR</sub> |
| --- | --- | --- | --- | --- | --- | --- | --- | --- |
| **BMI (kg/m<sup>2</sup>)** | 0.011 | 0.005 | 0.002 | 0.020 | 1698 | 2.333 | 0.020 | **0.020** |
| **Smoking (Current status)** | 0.106 | 0.044 | 0.019 | 0.192 | 1698 | 2.392 | 0.017 | **0.020** |
| **Education (years)** | -0.019 | 0.008 | -0.034 | -0.003 | 1698 | -2.394 | 0.017 | **0.020** |
| **Childhood trauma (CTQ Total)** | 0.044 | 0.017 | 0.010 | 0.078 | 2196 | 2.555 | 0.011 | **0.011** |
| **PRS for MDD** | 0.160 | 0.067 | 0.029 | 0.291 | 3901 | 2.395 | 0.017 | **0.025** |
| **PRS for BMI** | 0.041 | 0.019 | 0.004 | 0.077 | 3901 | 2.197 | 0.028 | **0.028** |
| **PRS for CRP** | 0.036 | 0.013 | 0.010 | 0.062 | 3901 | 2.666 | 0.008 | **0.023** |

After centering and scaling, each 1 standard deviation increase in the predictor is associated with a $\beta$ standard deviation increase in the brain age gap. PRS scores were residualized for 10 genetic PCs. Age, age<sup>2</sup>, and sex were added as covariates in the model. Random intercepts for scanning sites were included. FDR-corrected P-values considered statistically significant are indicated in **bold**.

### Association between the brain age gap and genetic factors

A higher brain age gap was associated with a higher PRS for BMI at one out of ten thresholds (β=0.029 SD increase per 1 SD [SE=0.013], p=0.026), higher PRS for CRP at six out of ten thresholds (β-values ranged from 0.01 to 0.045, P-values ranged from 0.001 to 0.015), and higher PRS for MDD at eight out of ten thresholds (β-values ranged from 0.023 to 0.038, P-values ranged from 0.003 to 0.026). An overview of these findings can be found in **Figure 1** and **Supplementary Tables S6-8.** There were no significant interactions between any of the PRS and diagnostic status (Control/MDD) on the brain age gap at any of the thresholds (**Supplementary Table S9**). When selecting the PRS at the threshold with the highest Z-value (categorical variable MDD) or t-value (continuous variables BMI and CRP) in the validation of PRS prediction and including them in one multivariable model, each 1 SD increase in genetic risk for MDD corresponded to a 0.16 SD increase in the brain age gap, while each 1 SD increase in genetic liability for elevated BMI or inflammation (CRP) were both associated with an 0.04 increase in the brain age gap (**Table 2**). Sensitivity analyses, which included the FreeSurfer version used for preprocessing as an additional covariate in the model, demonstrated that the association between the brain age gap and PRS for BMI became non-significant (β=0.035 SD per 1 SD [SE=0.018], p=0.055), suggesting that FreeSurfer version slightly moderates findings. All other findings remained unchanged.

**Figure 1.**
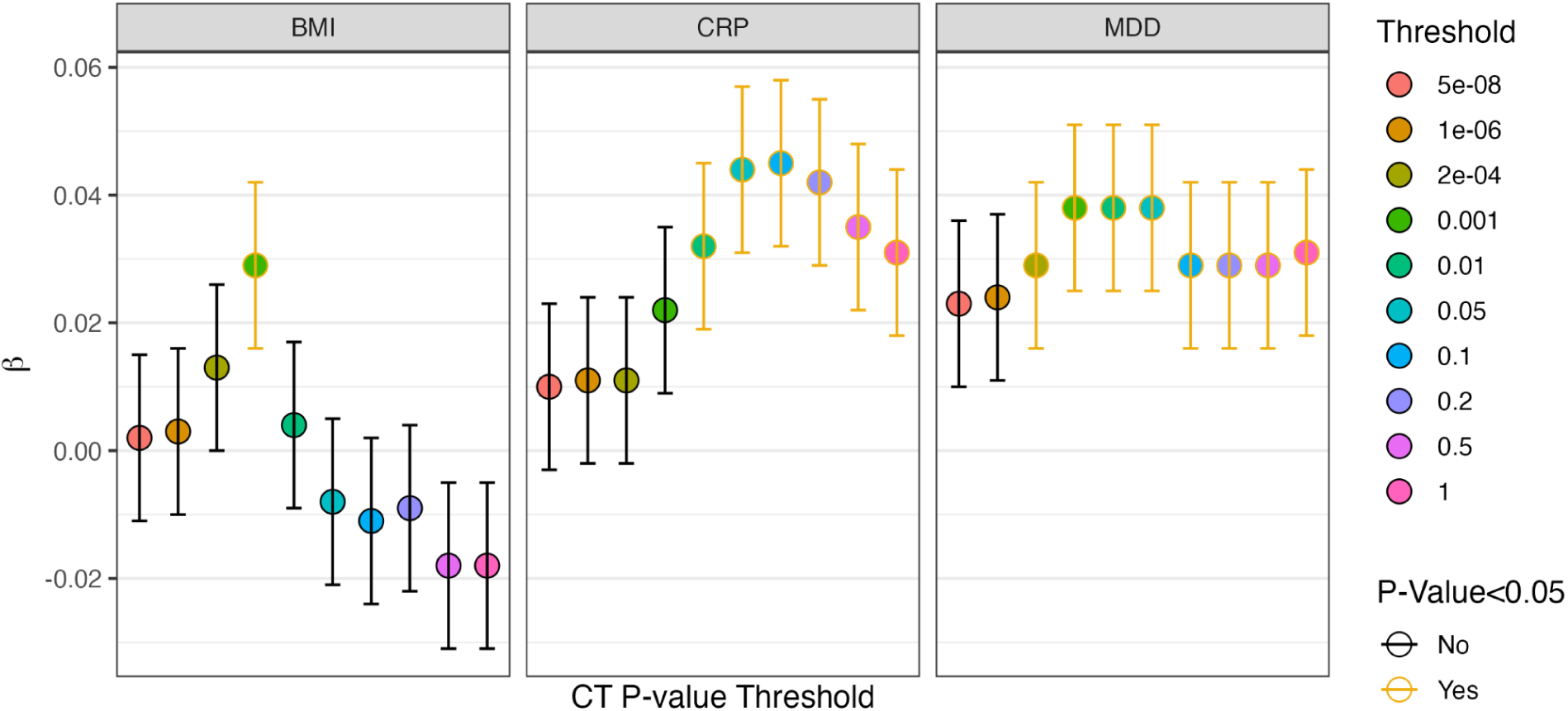
Associations between the brain age gap and genetic liability for body mass index, C-reactive protein, and major depressive disorder. The x-axis represents the PRS threshold. The y-axis represents the standardized regression coefficient. Abbreviations: BMI, body mass index; CRP, C-reactive protein; MDD, major depressive disorder; CT P-value Threshold, P-value threshold used for creating PRS using clumping and thresholding methods.

### Multivariable model

When all risk and resilience factors (lifestyle, early-life, genetic factors) were included in the same multivariable model, the associations between the brain age gap and childhood trauma and all three PRS became non-significant. However, BMI, current smoking status, and education remained uniquely and significantly associated with the brain age gap. In comparing the standardized regression coefficients of the different predictors within this multivariable model, smoking status exhibited the strongest association with the brain age gap (β = 0.113 SD per 1 SD, p=0.015), followed by BMI (β = 0.050 SD per 1 SD, p = 0.020) and education (β = -0.043 SD per 1 SD, p=0.032). These findings are summarized in **Table 3**. To evaluate whether the main effect of MDD diagnosis would be attenuated when accounting for lifestyle factors, we conducted a post-hoc supplementary analysis. This analysis showed that the effect of diagnosis became non-significant: individuals with MDD had, on average, a 0.04 SD higher brain age gap than controls (p = 0.26).

**Table 3.** Multivariable associations between the brain age gap and risk and resilience factors.

| Predictor | $\beta$ | lower CI | upper CI | SE | DF | t-value | P-value |
| --- | --- | --- | --- | --- | --- | --- | --- |
| **BMI (kg/m<sup>2</sup>)** | 0.050 | 0.008 | 0.092 | 0.021 | 1618 | 2.335 | **0.020** |
| **Smoking (Current status)** | 0.113 | 0.023 | 0.202 | 0.046 | 1618 | 2.446 | **0.015** |
| **Education (years)** | -0.043 | -0.082 | -0.004 | 0.02 | 1618 | -2.152 | **0.032** |
| **Childhood trauma (CTQ Total)** | 0.030 | -0.01 | 0.07 | 0.021 | 1618 | 1.447 | 0.148 |
| **PRS MDD** | -0.002 | -0.04 | 0.035 | 0.019 | 1618 | -0.114 | 0.909 |
| **PRS BMI** | 0.001 | -0.038 | 0.039 | 0.02 | 1618 | 0.033 | 0.974 |
| **PRS CRP** | 0.021 | -0.017 | 0.058 | 0.019 | 1618 | 1.074 | 0.283 |

After centering and scaling, each 1 standard deviation increase in the predictor is associated with a $\beta$ standard deviation increase in the brain age gap. PRS scores were residualized for 10 genetic PCs. Age, age<sup>2</sup>, and sex were added as covariates in the model. Random intercepts for scanning sites were included. P-values considered statistically significant are indicated in **bold**.

### Validation of PRS prediction of epigenetic clocks

For the validation of the PRS for GrimAge, PRS created at five out of ten thresholds were significantly associated with the GrimAge acceleration residuals with β-values ranging from 0.08 to 0.108 and P-values ranging from 0.002 to 0.034 (**Supplementary Table S10**). In the validation for the Hannum epigenetic clock, only PRS created at the 0.0002 threshold were significantly associated with Hannum age acceleration residuals (β=0.072, P=0.04; **Supplementary Table S11**). For the Horvath epigenetic clock, PRS at all ten thresholds were significantly associated with Horvath age acceleration residuals (β-values ranging from 0.077 to 0.169, P<0.029; **Supplementary Table S12**). Finally, for the PhenoAge epigenetic clock, PRS at eight out of ten thresholds were significantly associated with PhenoAge acceleration residuals, with β-values from 0.069 to 0.160 and P-values from 4.8×10^-6^ to 0.05 (**Supplementary Table S13**).

### Exploratory associations between the brain age gap and (PRS for) epigenetic clocks

Within the two cohorts with DNAm data available (N=810), we found that a higher brain age gap was associated with both “second-generation” epigenetic clocks, namely, the GrimAge acceleration residuals (β=0.139 years per year, P_FDR_<0.001) and PhenoAge acceleration residuals (β=0.070 years per year, P_FDR_ = 0.037), but not with the “first-generation” epigenetic clocks (i.e., Hannum and Horvath acceleration residuals; **Supplementary Table S14**). However, when analyzing the genetic liability for higher epigenetic age acceleration residuals across the entire genetic sample (N=3,934) in relation to the brain age gap, we only observed significant but distinct patterns with the “first-generation” epigenetic clocks. A higher brain age gap was associated with a lower genetic liability for epigenetic age acceleration as measured by Horvath acceleration residuals at two of ten thresholds (β-values ranging from -0.037 to -0.034 and P-values between 0.004 and 0.009), but with a higher genetic liability as measured by Hannum acceleration residuals at two of ten thresholds (β-values ranging from 0.028 to 0.030 and P-values between 0.032 and 0.020). An overview of these findings is displayed in **Figure 2** and **Supplementary Tables S15-18**.

**Figure 2.**
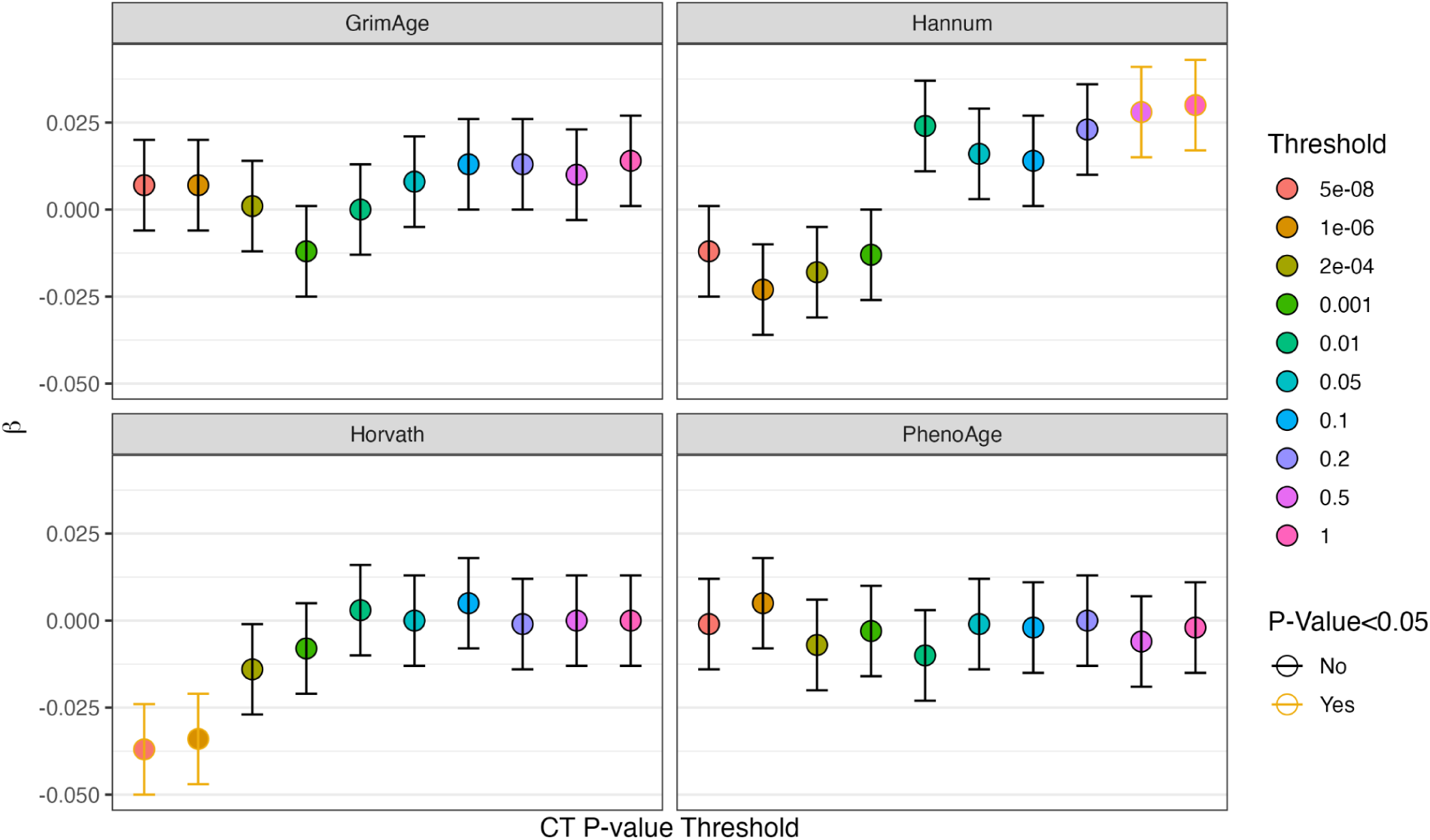
Associations between the brain age gap and genetic liability for epigenetic age acceleration residuals. The x-axis represents the PRS threshold. The y-axis represents the standardized regression coefficient. The GrimAge, Hannum, Horvath, and PhenoAge epigenetic clock panels indicate the PRS for the acceleration residuals. Abbreviations: CT P-value Threshold, P-value threshold used for creating PRS using clumping and thresholding methods.

## DISCUSSION

In this extensive multisite study, we pooled imaging, genetic, and phenotypic data from 12 international cohorts encompassing a total of N=3,934 individuals (N=1,846 controls and N=2,088 individuals with MDD). Our findings indicate that a higher genetic liability for major depression, as well as other depression-related traits (i.e., PRS for BMI and for CRP), significantly contributed to a higher brain age gap. This suggests that advanced brain age may partly stem from a (genetic predisposition to) depression and traits more commonly occurring in individuals with depression. Factors such as childhood trauma, smoking, and education also significantly contributed to the brain age gap, linking health risk factors to individual variations in age-related brain structure. When examined in a multivariable model, BMI, smoking, and education emerged as most uniquely and robustly associated with the brain age gap. These results imply that while genetic liability for depression and related traits is linked to poorer brain health, lifestyle factors potentially present opportunities for intervention. This interpretation is further supported by the post-hoc analysis, which showed that the previous significant difference in brain age gap between individuals with MDD and controls was attenuated when lifestyle factors were taken into account. Therefore, lifestyle interventions, including smoking cessation and weight management, may serve as (treatment) components that warrant stronger emphasis and implementation to prevent and treat brain and mental illness.

Several mechanisms may explain how the factors currently considered can impact the brain and can thereby affect behavior. First, emerging evidence on biological aging suggests that an unhealthy BMI contributes to cellular damage, leading to senescence and apoptosis, which in turn can trigger inflammation and reduced ability for repair.^57,58^ Neuroimmune mechanisms such as pro-inflammatory cytokines (e.g., CRP), affect biological processes like synaptic plasticity, and inflammatory biomarkers are commonly dysregulated in (subgroups of) MDD.^59^ As such, MDD represents a condition where inflammation, obesity, and premature or advanced aging co-occur and converge. Our findings support the role of genetic liability for elevated inflammation and BMI in linking the higher brain age gap observed in MDD. Second, to better understand the factors contributing to the brain age gap, we must also appreciate the structural brain features used to make accurate brain age predictions. Recent research has shown a causal effect of increased BMI on lower cortical thickness.^60^ Cortical thickness features were most important for the brain age algorithm to make accurate predictions (compared to surface area or subcortical volume). Notably, higher BMI, lower cortical thickness, and higher brain age gaps represent transdiagnostic patterns that are not exclusive to depression but also observed in other mental disorders such as bipolar disorder and schizophrenia.^61^ Additionally, about 18.4% of the total association between bipolar disorder and ventricular volume can be explained by a higher BMI.^62^ A qualitative comparison found that individuals with MDD with the highest decile of brain age gap values also had larger lateral ventricles compared to the bottom 90%.^22^ Taken together, these findings underscore the overlap between age-related and disease-related structural brain patterns. Finally, while perhaps unsurprising, we found no interaction between diagnosis and any of the health risk factors on the brain age gap. This aligns with existing evidence suggesting that a healthy lifestyle may not only help prevent depression,^63^ but also contributes to the maintenance of brain health.^33,64^ Overall, the current findings indicate that a higher brain age gap may arise from both increased genetic vulnerabilities to depression and related traits, as well as the consequences of having MDD. In this sense, the brain age gap could potentially serve as a clinical tool, providing an intuitive proxy for brain health that reflects a combination of genetic factors and the cumulative impact of health risk factors.

The present multivariable findings may inform the treatment of MDD, in that smoking and being overweight contributed more robustly and uniquely to the brain age gap than the genetic liabilities. While first-line psychotherapy and/or antidepressant medications are important components of MDD treatment, integrating lifestyle changes may further enhance both overall and immunometabolic outcomes.^65–67^ This approach could serve as a particularly valuable alternative for subgroups of individuals with MDD or other mental health disorders who are impacted by psychotropic drug-related weight gain, obesity, and/or associated cardiometabolic comorbidities.^68,69^ Our findings thus offer a neurobiological rationale for placing a stronger emphasis on involving lifestyle components, such as smoking cessation and weight management, which could significantly benefit both mental and brain health.^70,71^ Even so, the current findings are based on pooled observational studies, and their clinical implications should be interpreted with caution. To date, the prognostic value of the brain age gap on (antidepressant) treatment outcomes is scarce and inconsistent.^26,72^ Future research could investigate whether the brain age gap might inform alternative treatment strategies. For instance, it would be valuable to test whether combining GLP-1 receptor agonists or anti-inflammatory agents with lifestyle interventions could yield specific benefits for depressed individuals with a higher brain age gap. Consequently, well-controlled studies are crucial to demonstrate whether the brain age gap can be useful in a clinical scenario.

MDD is not only linked to a higher brain age gap,^25^ but also to several first-^73^ and second-generation^74,75^ epigenetic clocks in both young and adult populations.^8,76–78^ A recent Mendelian randomization study revealed that depression liability was potentially causally related to higher GrimAge acceleration residuals (i.e., indicating advanced epigenetic age).^79^ In our exploratory analyses involving two cohorts with available DNAm data (N=810), we found that each additional year in the GrimAge and PhenoAge acceleration residuals predicted a 51-day and 26-day increase in the brain age gap, respectively. However, two out of four epigenetic clocks (the “first-generation” clocks) showed no significant relationship with the brain age gap. These findings align with other studies demonstrating that the brain age gap is largely independent of, or only weakly correlated with, epigenetic clocks in late adolescence,^80^ as well as in adult^81^ and elderly samples.^82^ Overall, there appears to be a low agreement between imaging-derived age predictors and those derived from peripheral measurements in general,^23^ and specifically with DNAm-derived age predictors. This discrepancy raises intriguing questions about the differences in central versus peripheral aging, however, such interpretations about an ongoing process of aging may not be appropriate without supporting longitudinal data.^83^ Nevertheless, a significant advantage is that different biological age indicators can thus be combined to produce stronger effect sizes for health determinants^9^ and more accurate predictions of global cognitive function status^84^ and mortality.^82^

### Strengths and limitations

A key strength of the current study is the use of standardized protocols for both brain age and PRS calculations, which minimizes analytic heterogeneity and enhances the generalizability of our findings across 12 cohorts worldwide. In addition, large sample sizes are needed to find reliable links between brain imaging phenotypes, genetics, and behavior.^85^ Another advantage is that our outcome measure, the brain age gap (which condenses complex patterns of age-related brain patterns into a single score), and our genetic predictor variables (PRS, summarizing the widespread genetic risk for depression and related traits into a single score) optimize statistical power. By greatly reducing the number of comparisons between imaging and genetics, this approach improves the precision of brain-genetic-behavior correlations and provides valuable insights into the contributions to individual differences in the brain age gap. However, the generalizability of our findings is also limited by certain factors. Both the training data for brain age prediction models and the discovery samples used for GWAS to derive the PRS are primarily composed of individuals of European descent. Previous studies have shown ethnic differences in imaging-derived phenotypes across diverse populations,^86–88^ suggesting that genetic ancestry influences brain structure. Such genetic differences may affect imaging phenotypes and contribute to varying risks for conditions such as depression.^89,90^ To improve our understanding of brain-genetic-behavior interactions across ethnic and geographic boundaries, future research should prioritize the inclusion of globally diverse populations, for example by extending to Chinese samples through international collaborations with the Depression Imaging REsearch ConsorTium (DIRECT) consortium.^91^ A natural progression of this study would also be to examine a broader set of lifestyle and health risk factors - such as diet, air pollution, socioeconomic background, and urbanization - that are associated with individual differences in brain structure ^92,93^ but were not addressed in this study. Finally, future research should test other methods and modelling advancements that further improve brain age reliability. For example, by incorporating uncertainty estimation in brain age models, to enhance out-of-distribution detection, transparency, and interpretability.^94^

### Conclusions

The findings of this observational study suggest that modifiable health behaviors, such as smoking and BMI, contribute to poorer brain health, extending beyond the genetic predispositions to adverse depression and metabolic profiles. Therefore, further longitudinal research is needed to determine whether smoking cessation and weight management could be effective in enhancing brain health, especially within the context of clinical trials. This research is crucial to confirm whether lifestyle interventions—perhaps currently underutilized in depression treatment—can improve both mental and brain health. Integrated mental and physical healthcare approaches in psychiatry may promote significant (brain) health gains and cost savings, by prevention, earlier identification, management and treatment.

## Conflicts of Interest Disclosures

HJG has received travel grants and speakers honoraria from Neuraxpharm, Servier, Indorsia and Janssen Cilag.

IH holds a 3.2% equity share in Innowell Pty Ltd that is focused on digital transformation of mental health services.

JHC is scientific advisor to and shareholder in Brain Key and Claritas HealthTech PTE.

YM has received consulting fees from Noema Pharma not related to the present study. All other authors do not report any conflicts of interest.

## Acknowledgments

Consortium and cohorts

- *ENIGMA MDD* is supported by NIH (R01 MH129832 Morey & Schmaal; R01 MH129742 Thompson).
- The *BiDirect* Study was funded by grants of the German Federal Ministry of Education and Research (BMBF; Grants FKZ-01ER0816 and FKZ-01ER1506) to Klaus Berger.
- The *FOR2107* cohorts were funded by the German Research Foundation (DFG grants FOR2107 KI588/14-1, and KI588/14-2, and KI588/20-1, KI588/22-1 to Tilo Kircher, Marburg, Germany). Biosamples and corresponding data were sampled, processed and stored in the Marburg Biobank CBBMR. This work was also funded by the German Research Foundation (DFG, grant FOR2107 DA1151/5-1, DA1151/5-2, DA1151/9-1, DA1151/10-1, DA1151/11-1 to UD; SFB/TRR 393, project grant no 521379614) and the Interdisciplinary Center for Clinical Research (IZKF) of the medical faculty of Münster (grant Dan3/022/22 to UD).
- The *MOTAR* study (www.motar.nl) has been funded through the Netherlands Organisation for Health Research and Development (VICI grant number 91811602) and funds from VU University Medical Center and GGZ inGeest.
- The *MPIP* Sample comprises patients and control subjects included in the Munich Antidepressant Response Signature study and the Recurrent Unipolar Depression (RUD) Case-Control study. We wish to acknowledge Rosa Schirmer, Elke Schreiter, Reinhold Borschke and Ines Eidner for image acquisition and data preparation, Anna Olynyik for data quality control, and Benno Pütz, Nazanin Karbalai, and Bertram Müller-Myhsok for distributed computing support. We thank Darina Czamara and Xinyang Yu for supporting the quality control of the epigenetic data. We thank Dorothee P. Auer and Florian Holsboer for initiation of the RUD study. The MARS study was supported by a grant of the Exzellenz-Stiftung of the Max Planck Society. This work has also been funded by the BMBF in the framework of the National Genome Research Network (NGFN) (FKZ 01GS0481). The funders had no role in study design, data collection and analysis, decision to publish, or preparation of the manuscript.
- This work was additionally funded by the “Innovative Medizinische Forschung” (IMF) by the medical faculty of the University of Münster awarded to SM (ME122205), the Else Kröner-Fresenius Stiftung awarded to SM (2023_EKEA.153). This work was supported in part by the consortium grant Trajectories of Affective Disorders from the German Research Foundation (DFG) SFB/TRR 393 (project grant no 521379614).
- The infrastructure for the *NESDA study* (www.nesda.nl) is funded through the Geestkracht program of the Netherlands Organisation for Health Research and Development (ZonMw, grant number 10-000-1002) and financial contributions by participating universities and mental health care organizations (VU University Medical Center, GGZ inGeest, Leiden University Medical Center, Leiden University, GGZ Rivierduinen, University Medical Center Groningen, University of Groningen, Lentis, GGZ Friesland, GGZ Drenthe, Rob Giel Onderzoekscentrum).
- *SHIP* is part of the Community Medicine Research net of the University of Greifswald, Germany, which is funded by the Federal Ministry of Education and Research (grants no. 01ZZ9603, 01ZZ0103, and 01ZZ0403), the Ministry of Cultural Affairs and the Social Ministry of the Federal State of Mecklenburg-West Pomerania. Genome-wide SNP typing in *SHIP* and MRI scans in *SHIP-Start* and *SHIP-TREND-0* have been supported by a joint grant from Siemens Healthineers, Erlangen, Germany and the Federal State of Mecklenburg-West Pomerania.
- The ENIGMA consortium received funding from the National Institutes of Health (NIH) Consortium grant U54 EB020403, R01MH131806, R01MH129742, and is supported by a cross-NIH alliance that funds Big Data to Knowledge Centers of Excellence (BD2K).

*Authors*

- AP is supported by a scholarship/stipend from Deutscher Akademischer Austauschdienst (DAAD).
- LH was funded by the Rubicon (grant number 452020227) and Veni award (grant number 09150162210201) from the Dutch Research Council (NWO).
- LS is supported by an NHMRC Investigator Leadership Grant (2017962) and a University of Melbourne Dame Kate Campbell Fellowship.
- CML. was supported by the Heisenberg programme of the German Research Research Foundation (DFG; LI 2654/4-1). This work was partly funded by the EU Joint Programme – Neurodegenerative Disease Research (JPND2021-650-289, coordinator: CML).
- IH is supported by an NHMRC Research Fellowship GNT2016346 2023-2027: Right care, first time: delivering technology-enabled mental health care to young people at scale.
- NJ is supported by R01MH134004 from the NIH.
- SEM is supported by NHMRC APP2025674.
- SIT was supported by R01MH116147, R01AG058854, R01MH129742.
- YM is partially supported by Amsterdam UMC (Starter Grant Ronde 2) and Amsterdam Neuroscience (PoC funding 2024-2026).
- HW is supported by the Wellcome Trust Strategic Award (104036/Z/14/Z)

## Data Sharing Statement

The datasets analyzed during the current study are not publicly available due to site restrictions but data may be available from the corresponding sites on reasonable request.

## REFERENCES

1. Scott KM, Lim C, Al-Hamzawi A, et al. Association of mental disorders with subsequent chronic physical conditions: World mental health surveys from 17 countries. JAMA Psychiatry. 2016;73(2):150–158.

2. Révész D, Verhoeven JE, Milaneschi Y, Penninx BWJH. Depressive and anxiety disorders and short leukocyte telomere length: mediating effects of metabolic stress and lifestyle factors. Psychol Med. 2016;46(11):2337–2349.

3. Darrow SM, Verhoeven JE, Révész D, et al. The Association Between Psychiatric Disorders and Telomere Length: A Meta-Analysis Involving 14,827 Persons. Psychosom Med. 2016;78(7):776–787.

4. Penninx BWJH. Depression and cardiovascular disease: Epidemiological evidence on their linking mechanisms. Neurosci Biobehav Rev. Published online 2016. doi:10.1016/j.neubiorev.2016.07.003

5. Pan A, Keum N, Okereke OI, et al. Bidirectional association between depression and metabolic syndrome: a systematic review and meta-analysis of epidemiological studies. Diabetes Care. 2012;35(5):1171–1180.

6. Vigo D, Thornicroft G, Atun R. Estimating the true global burden of mental illness. Lancet Psychiatry. 2016;3(2):171–178.

7. Marx W, Penninx BWJH, Solmi M, et al. Major depressive disorder. Nat Rev Dis Primers. 2023;9(1):44.

8. Han LKM, Aghajani M, Penninx BWJH, Copeland WE, Aberg KA, van den Oord EJCG. Lagged effects of childhood depressive symptoms on adult epigenetic aging. Psychol Med. Published online October 7, 2024:1–9.

9. Jansen R, Han LK, Verhoeven JE, et al. An integrative study of five biological clocks in somatic and mental health. Elife. 2021;10. doi:10.7554/eLife.59479

10. Firth J, Siddiqi N, Koyanagi A, et al. The Lancet Psychiatry Commission: a blueprint for protecting physical health in people with mental illness. Lancet Psychiatry. 2019;6(8):675–712.

11. Penninx BWJH, Lange SMM. Metabolic syndrome in psychiatric patients: overview, mechanisms, and implications. Dialogues Clin Neurosci. 2018;20(1):63–73.

12. Smith DJ, Court H, McLean G, et al. Depression and multimorbidity: a cross-sectional study of 1,751,841 patients in primary care: A cross-sectional study of 1,751,841 patients in primary care. J Clin Psychiatry. 2014;75(11):1202–1208; quiz 1208.

13. Simon J, Wienand D, Park AL, et al. Excess resource use and costs of physical comorbidities in individuals with mental health disorders: A systematic literature review and meta-analysis. Eur Neuropsychopharmacol. 2023;66:14–27.

14. Han LKM, Verhoeven JE, Tyrka AR, et al. Accelerating research on biological aging and mental health: Current challenges and future directions. Psychoneuroendocrinology. 2019;106:293–311.

15. Epel ES, Prather AA. Stress, Telomeres, and Psychopathology: Toward a Deeper Understanding of a Triad of Early Aging. Annu Rev Clin Psychol. 2018;14:371–397.

16. Lindqvist D, Epel ES, Mellon SH, et al. Psychiatric disorders and leukocyte telomere length: Underlying mechanisms linking mental illness with cellular aging. Neurosci Biobehav Rev. 2015;55:333–364.

17. Verhoeven JE, Révész D, Epel ES, Lin J, Wolkowitz OM, Penninx BWJH. Major depressive disorder and accelerated cellular aging: results from a large psychiatric cohort study. Mol Psychiatry. 2013;(May):1–7.

18. Walker ER, McGee RE, Druss BG. Mortality in mental disorders and global disease burden implications: a systematic review and meta-analysis. JAMA Psychiatry. 2015;72(4):334–341.

19. Cole JH, Franke K. Predicting Age Using Neuroimaging: Innovative Brain Ageing Biomarkers. Trends Neurosci. 2017;40(12):681–690.

20. Gaser C, Franke K. 10 years of BrainAGE as an neuroimaging biomarker of brain aging: What insights did we gain? Front Neurol. 2019;10:789.

21. Cole JH, Marioni RE, Harris SE, Deary IJ, Cole JH. Brain age and other bodily “ ages ” : implications for neuropsychiatry. Mol Psychiatry. Published online 2018. doi:10.1038/s41380-018-0098-1

22. Han LKM, Dinga R, Hahn T, et al. Brain aging in major depressive disorder: results from the ENIGMA major depressive disorder working group. Mol Psychiatry. Published online May 18, 2020. doi:10.1038/s41380-020-0754-0

23. Han LKM, Schnack HG, Brouwer RM, et al. Contributing factors to advanced brain aging in depression and anxiety disorders. Transl Psychiatry. 2021;11(1):402.

24. Han LKM, Dinga R, Leenings R, Hahn T, Cole JH. A large-scale ENIGMA multisite replication study of brain age in depression. Neuroimage. Published online 2022. https://www.sciencedirect.com/science/article/pii/S2666956022000733

25. Ballester PL, Romano MT, de Azevedo Cardoso T, et al. Brain age in mood and psychotic disorders: a systematic review and meta-analysis. Acta Psychiatr Scand. 2022;145(1):42–55.

26. Ballester PL, Suh JS, Nogovitsyn N, et al. Accelerated brain aging in major depressive disorder and antidepressant treatment response: A CAN-BIND report. Neuroimage Clin. 2021;32:102864.

27. Dunlop K, Victoria LW, Downar J, Gunning FM, Liston C. Accelerated brain aging predicts impulsivity and symptom severity in depression. Neuropsychopharmacology. 2021;46(5):911–919.

28. Schmaal L, Pozzi E, C Ho T, et al. ENIGMA MDD: seven years of global neuroimaging studies of major depression through worldwide data sharing. Transl Psychiatry. 2020;10(1):172.

29. Wrigglesworth J, Ward P, Harding IH, et al. Factors associated with brain ageing - a systematic review. BMC Neurol. 2021;21(1):312.

30. Mouches P, Wilms M, Bannister JJ, Aulakh A, Langner S, Forkert ND. An exploratory causal analysis of the relationships between the brain age gap and cardiovascular risk factors. Front Aging Neurosci. 2022;14:941864.

31. Steffener J, Habeck C, O’Shea D, Razlighi Q, Bherer L, Stern Y. Differences between chronological and brain age are related to education and self-reported physical activity. Neurobiol Aging. 2016;40(February):138–144.

32. Jawinski P, Markett S, Drewelies J, et al. Linking brain age gap to mental and physical health in the Berlin Aging Study II. Front Aging Neurosci. 2022;14:791222.

33. Bittner N, Jockwitz C, Franke K, et al. When your brain looks older than expected: combined lifestyle risk and BrainAGE. Brain Struct Funct. 2021;226(3):621–645.

34. Cole JH. Multimodality neuroimaging brain-age in UK biobank: relationship to biomedical, lifestyle, and cognitive factors. Neurobiol Aging. 2020;92:34–42.

35. Pang Y, Zhao S, Zhang Z, et al. Individual structural covariance connectome reveals aberrant brain developmental trajectories associated with childhood maltreatment. J Psychiatr Res. 2025;181:709–715.

36. Beck D, Whitmore L, MacSweeney N, et al. Dimensions of early-life adversity are differentially associated with patterns of delayed and accelerated brain maturation. Biol Psychiatry. 2025;97(1):64–72.

37. Keding TJ, Heyn SA, Russell JD, et al. Differential patterns of delayed emotion circuit maturation in abused girls with and without internalizing psychopathology. Am J Psychiatry. 2021;178(11):1026–1036.

38. Choi SW, Mak TSH, O’Reilly PF. Tutorial: a guide to performing polygenic risk score analyses. Nat Protoc. 2020;15(9):2759–2772.

39. Kaufmann T, van der Meer D, Doan NT, et al. Common brain disorders are associated with heritable patterns of apparent aging of the brain. Nat Neurosci. 2019;22(10):1617–1623.

40. Yi F, Yuan J, Somekh J, et al. Genetically supported targets and drug repurposing for brain aging: A systematic study in the UK Biobank. Sci Adv. 2025;11(11):eadr3757.

41. Leonardsen EH, Peng H, Kaufmann T, et al. Deep neural networks learn general and clinically relevant representations of the ageing brain. Neuroimage. 2022;256(119210):119210.

42. Seitz-Holland J, Haas SS, Penzel N, Reichenberg A, Pasternak O. BrainAGE, brain health, and mental disorders: A systematic review. Neurosci Biobehav Rev. 2024;159(105581):105581.

43. Horvath S. DNA methylation age of human tissues and cell types. Genome Biol. 2013;14(10):R115.

44. Hannum G, Guinney J, Zhao L, et al. Genome-wide Methylation Profiles Reveal Quantitative Views of Human Aging Rates. Mol Cell. 2013;49(2):359–367.

45. Lu AT, Quach A, Wilson JG, et al. DNA methylation GrimAge strongly predicts lifespan and healthspan. Aging. 2019;11(2):303.

46. Levine ME, Lu AT, Quach A, et al. An epigenetic biomarker of aging for lifespan and healthspan. Aging. 2018;10(4):573–591.

47. Fischl B. FreeSurfer. Neuroimag*e2*. 2012;62(2):774–781.

48. Howard DM, Adams MJ, Clarke TK, et al. Genome-wide meta-analysis of depression identifies 102 independent variants and highlights the importance of the prefrontal brain regions. Nat Neurosci. 2019;22(3):343.

49. Locke AE, Kahali B, Berndt SI, et al. Genetic studies of body mass index yield new insights for obesity biology. Nature. 2015;518(7538):197–206.

50. Ligthart S, Vaez A, Võsa U, et al. Genome analyses of> 200,000 individuals identify 58 loci for chronic inflammation and highlight pathways that link inflammation and complex disorders. Am J Hum Genet. 2018;103(5):691–706.

51. Turchin MC, Hirschhorn JN. Gencrypt: one-way cryptographic hashes to detect overlapping individuals across samples. Bioinformatics. 2012;28(6):886–888.

52. Choi SW, O’Reilly PF. PRSice-2: Polygenic Risk Score software for biobank-scale data. Gigascience. 2019;8(7). doi:10.1093/gigascience/giz082

53. Yang J, Lee SH, Goddard ME, Visscher PM. GCTA: a tool for genome-wide complex trait analysis. Am J Hum Genet. 2011;88(1):76–82.

54. Bernstein DP, Fink L, Handelsman L, et al. Initial reliability and validity of a new retrospective measure of child abuse and neglect. Am J Psychiatry. 1994;151(8):1132–1136.

55. Bernstein DP, Stein JA, Newcomb MD, et al. Development and validation of a brief screening version of the Childhood Trauma Questionnaire. Child Abuse Negl. 2003;27(2):169–190.

56. McCartney DL, Min JL, Richmond RC, et al. Genome-wide association studies identify 137 genetic loci for DNA methylation biomarkers of aging. Genome Biol. 2021;22(1):194.

57. López-Otín C, Blasco MA, Partridge L, Serrano M, Kroemer G. Hallmarks of aging: An expanding universe. Cell. 2023;186(2):243–278.

58. Milaneschi Y, Simmons WK, van Rossum EFC, Penninx BW. Depression and obesity: evidence of shared biological mechanisms. Mol Psychiatry. 2019;24(1):18–33.

59. Frank P, Jokela M, Batty GD, Cadar D, Steptoe A, Kivimäki M. Association between systemic inflammation and individual symptoms of depression: A pooled analysis of 15 population-based cohort studies. Am J Psychiatry. 2021;178(12):1107–1118.

60. Opel N, Painter J, Refisch A, et al. Deciphering the Causal Influence of BMI and related Metabolic, Inflammatory, and Cardiovascular Factors on Brain Structure: A Mendelian Randomization Study. Published online 2024. https://www.researchsquare.com/article/rs-4365189/latest

61. Constantinides C, Han LKM, Alloza C, et al. Brain ageing in schizophrenia: evidence from 26 international cohorts via the ENIGMA Schizophrenia consortium. bioRxiv. Published online January 11, 2022. doi:10.1101/2022.01.10.21267840

62. McWhinney SR, Abé C, Alda M, et al. Association between body mass index and subcortical brain volumes in bipolar disorders-ENIGMA study in 2735 individuals. Mol Psychiatry. 2021;26(11):6806–6819.

63. Zhao Y, Yang L, Sahakian BJ, et al. The brain structure, immunometabolic and genetic mechanisms underlying the association between lifestyle and depression. Nat Ment Health. 2023;1(10):736–750.

64. Dhana K, Agarwal P, James BD, et al. Healthy lifestyle and cognition in older adults with common neuropathologies of dementia. JAMA Neurol. 2024;81(3):233–239.

65. Verhoeven JE, Han LKM, Lever-van Milligen BA, et al. Antidepressants or running therapy: Comparing effects on mental and physical health in patients with depression and anxiety disorders. J Affect Disord. 2023;329:19–29.

66. Lopresti AL. It is time to investigate integrative approaches to enhance treatment outcomes for depression? Medical Hypotheses. 2019;126:82–94.

67. Vreijling SR, Penninx BWJH, Verhoeven JE, et al. Running therapy or antidepressants as treatments for immunometabolic depression in patients with depressive and anxiety Disorders: A secondary analysis of the MOTAR study. Brain Behav Immun. 2024;123:876–883.

68. McIntyre RS, Kwan ATH, Rosenblat JD, Teopiz KM, Mansur RB. Psychotropic drug-related weight gain and its treatment. Am J Psychiatry. 2024;181(1):26–38.

69. Milaneschi Y, Lamers F, Berk M, Penninx BWJH. Depression heterogeneity and its biological underpinnings: Toward immunometabolic depression. Biol Psychiatry. 2020;88(5):369–380.

70. Marx W, Manger SH, Blencowe M, et al. Clinical guidelines for the use of lifestyle-based mental health care in major depressive disorder: World Federation of Societies for Biological Psychiatry (WFSBP) and Australasian Society of Lifestyle Medicine (ASLM) taskforce. World J Biol Psychiatry. 2023;24(5):333–386.

71. Sarris J, O’Neil A, Coulson CE, Schweitzer I, Berk M. Lifestyle medicine for depression. BMC Psychiatry. 2014;14(1):107.

72. Jha MK, Fatt CC, Minhajuddin A, Mayes T, Trivedi MH. Accelerated brain aging in adults with major depressive disorder predicts poorer outcome with sertraline: Findings from the EMBARC study. Biol Psychiatry Cogn Neurosci Neuroimaging. Published online September 27, 2022. doi:10.1016/j.bpsc.2022.09.006

73. Liu C, Wang Z, Hui Q, et al. Association between depression and epigenetic age acceleration: A co-twin control study. Depress Anxiety. 2022;39(12):741–750.

74. Protsenko E, Yang R, Nier B, et al. “GrimAge,” an epigenetic predictor of mortality, is accelerated in major depressive disorder. Transl Psychiatry. 2021;11(1):193.

75. Rampersaud R, Protsenko E, Yang R, et al. Dimensions of childhood adversity differentially affect biological aging in major depression. Transl Psychiatry. 2022;12(1):431.

76. Han LKM, Aghajani M, Clark SL, et al. Epigenetic Aging in Major Depressive Disorder. Am J Psychiatry. 2018;175(8):774–782.

77. Li Z, He Y, Ma X, Chen X. Epigenetic age analysis of brain in major depressive disorder. Psychiatry Res. 2018;269:621–624.

78. Shindo R, Tanifuji T, Okazaki S, et al. Accelerated epigenetic aging and decreased natural killer cells based on DNA methylation in patients with untreated major depressive disorder. NPJ Aging. 2023;9(1):19.

79. Luo X, Ruan Z, Liu L. Causal relationship between depression and aging: a bidirectional two-sample Mendelian randomization study. Aging Clin Exp Res. 2023;35(12):3179–3187.

80. Sanders F, Baltramonaityte V, Donohoe G, et al. Associations between methylation age and brain age in late adolescence. bioRxiv. Published online September 10, 2022. doi:10.1101/2022.09.08.506972

81. Teeuw J, Ori APS, Brouwer RM, et al. Accelerated aging in the brain, epigenetic aging in blood, and polygenic risk for schizophrenia. Schizophr Res. 2021;231:189–197.

82. Cole JH, Ritchie SJ, Bastin ME, et al. Brain age predicts mortality. Mol Psychiatry. 2018;23(5):1385–1392.

83. Ikram MA. The use and misuse of “biological aging” in health research. Nat Med. Published online October 7, 2024. doi:10.1038/s41591-024-03297-9

84. Zheng Y, Habes M, Gonzales M, et al. Mid-life epigenetic age, neuroimaging brain age, and cognitive function: coronary artery risk development in young adults (CARDIA) study. Aging. 2022;14(4):1691–1712.

85. Marek S, Tervo-Clemmens B, Calabro FJ, et al. Reproducible brain-wide association studies require thousands of individuals. Nature. 2022;603(7902):654–660.

86. Chee MWL, Zheng H, Goh JOS, Park D, Sutton BP. Brain structure in young and old East Asians and Westerners: comparisons of structural volume and cortical thickness. J Cogn Neurosci. 2011;23(5):1065–1079.

87. Long H, Liu B, Hou B, et al. The long rather than the short allele of 5-HTTLPR predisposes Han Chinese to anxiety and reduced connectivity between prefrontal cortex and amygdala. Neurosci Bull. 2013;29(1):4–15.

88. Tang Y, Zhao L, Lou Y, et al. Brain structure differences between Chinese and Caucasian cohorts: A comprehensive morphometry study. Hum Brain Mapp. 2018;39(5):2147–2155.

89. Bigdeli TB, Ripke S, Peterson RE, et al. Genetic effects influencing risk for major depressive disorder in China and Europe. Transl Psychiatry. 2017;7(3):e1074.

90. Meng X, Navoly G, Giannakopoulou O, et al. Multi-ancestry genome-wide association study of major depression aids locus discovery, fine mapping, gene prioritization and causal inference. Nat Genet. 2024;56(2):222–233.

91. Chen X, Lu B, Li HX, et al. The DIRECT consortium and the REST-meta-MDD project: towards neuroimaging biomarkers of major depressive disorder. Psychoradiology. 2022;2(1):32–42.

92. Liu F, Xu J, Guo L, et al. Environmental neuroscience linking exposome to brain structure and function underlying cognition and behavior. Mol Psychiatry. 2023;28(1):17–27.

93. Lin F, Chen X, Cai Y, et al. Accelerated biological aging as potential mediator mediates the relationship between pro-inflammatory diets and the risk of depression and anxiety: A prospective analysis from the UK biobank. J Affect Disord. 2024;355:1–11.

94. Hahn T, Ernsting J, Winter NR, et al. An uncertainty-aware, shareable, and transparent neural network architecture for brain-age modeling. Sci Adv. 2022;8(1):eabg9471.

