## Supplement for "Lifestyle, Early-life, and Genetic Health Risk Factors Underlying the Brain Age Gap: A Mega-Analysis Across 3,934 Individuals from the ENIGMA MDD Consortium"

### SUPPLEMENTARY FIGURES

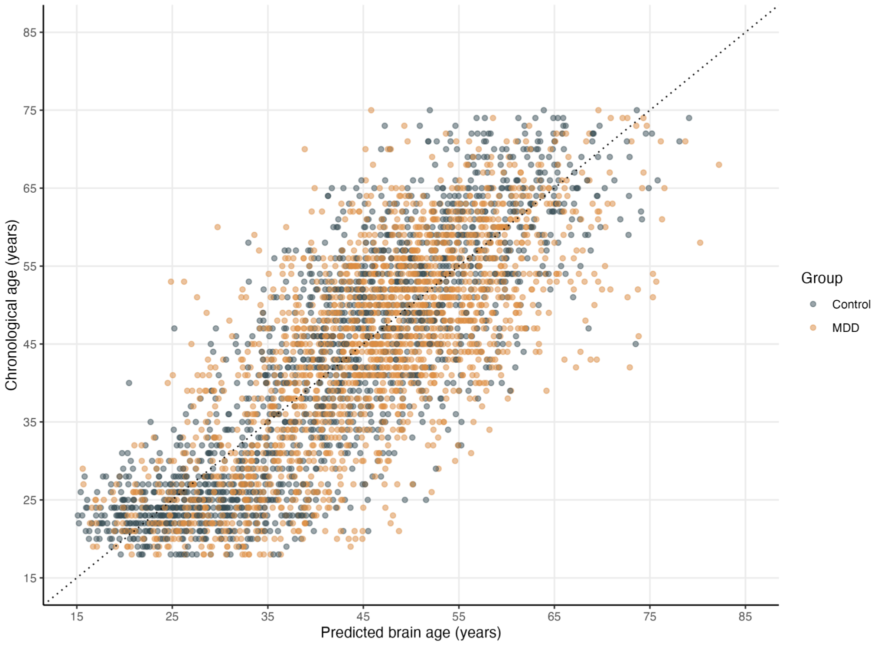


**Supplementary Figure S1. Predicted brain age against chronological age.**

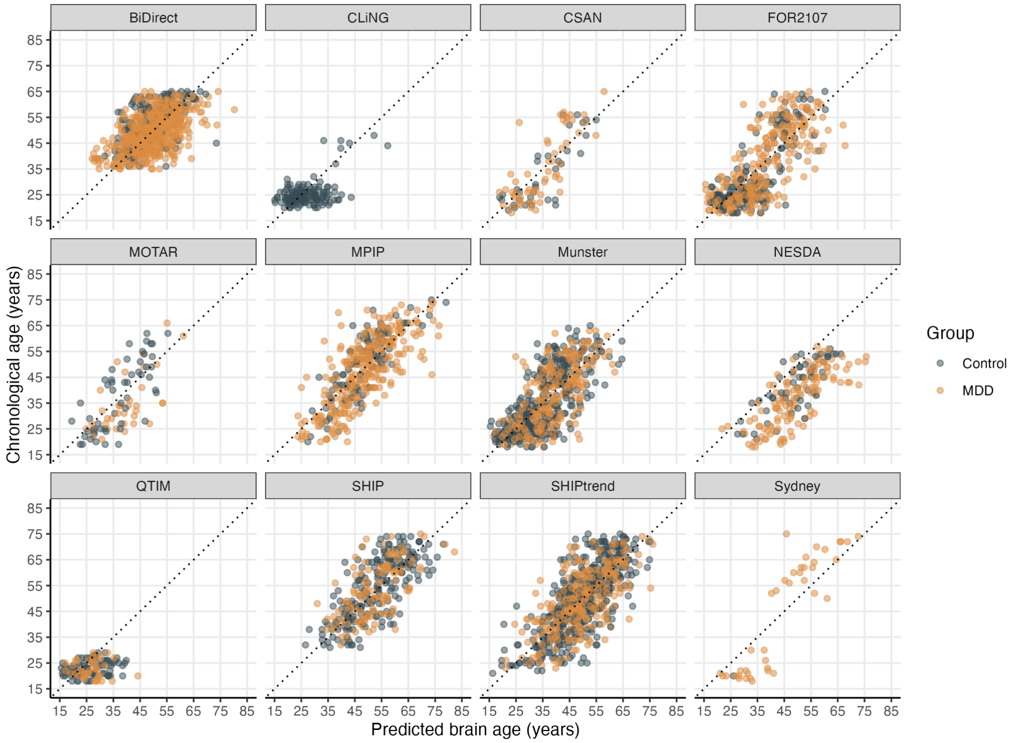


**Supplementary Figure S2. Predicted brain age against chronological age per cohort.**

### SUPPLEMENTARY TABLES

**Supplementary Table S1.** Cohort-specific details on diagnostic measurement and in- and exclusion criteria.

| Cohort | Country | Diagnosis measurement | Sample characteristics/Inclusion criteria | Exclusion criteria |
|---|---|---|---|---|
| **BiDirect** | Germany | M.I.N.I. Neuropsychiatric Interview, IDS, HAMD, CESD, ICD-10 | Patients hospitalized for a first or recurrent episode of depression, population controls randomly selected in city registry | dementia, addiction |
| **CSAN** | | M.I.N.I. Neuropsychiatric Interview | Current MDD: Meets MINI criteria for depression; comorbid anxiety disorders are allowed; mood-congruent psychotic symptoms allowed. | Current MDD: a current DSM-5 diagnosis of substance use disorder, except nicotine; a psychotic disorder, except depression with mood-congruent psychotic features; new antidepressant medication during the month before study participation (two months for fluoxetine); change of the dose of psychotropic medications over the last month (antidepressant and antipsychotic medication) or the last two months (mood stabilizers and anticonvulsants). |
| **CliNG** | Germany | ICD-10 interview | Patients met the diagnostic criteria for major depressive disorder according to ICD-10 classification standards and were aged between 18 and 60 years. | Exclusion criteria for MDD subjects were neurological and severe other medical conditions (in particular those that could be related to affective symptoms), lifetime diagnosis of substance dependence, substance abuse during the last month, cannabis abuse during the last 2 weeks, mental retardation as well as past or actual presence of other axis I diagnoses with exception of anxiety disorders. Exclusion criteria for control subjects were neurological, psychiatric and severe other medical conditions, lifetime diagnosis of substance dependence, substance abuse during the last month, cannabis abuse during the last 2 weeks, previous or actual use of psychotropic medication, and mental retardation. |

| | | | | |
|---|---|---|---|---|
| **FOR2107 - Marburg** | Germany | SCID-1 | Participants recruited by means of public advertisement and from the inpatient services. Inclusion criteria: age 18-65 years; patients were diagnosed with major depressive disorder by SCID-Interview, currently depressed or remitted. | Exclusion criteria all: any MRI contraindications; any neurological abnormalities. Exclusion criteria controls: any current or former psychiatric disorder; Exclusion criteria patients: substance dependence or current benzodiazepine treatment (wash out of at least three half-lives before study participation)" |
| **FOR2017 - Münster** | Germany | SCID-1 | Participants recruited by means of public advertisement and from the inpatient services. Inclusion criteria: age 18-65 years; patients were diagnosed with major depressive disorder by SCID-Interview, currently depressed or remitted. | Exclusion criteria all: any MRI contraindications; any neurological abnormalities. Exclusion criteria controls: any current or former psychiatric disorder; Exclusion criteria patients: substance dependence or current benzodiazepine treatment (wash out of at least three half-lives before study participation)" |
| **MOTAR** | Netherlands | CIDI interview | Inclusion criteria of the patient sample include: having a current depressive disorder (major depressive disorder) or anxiety disorder (social phobia, generalized anxiety disorder, panic disorder with or without agoraphobia) as ascertained by the Diagnostic and Statistical Manual of Mental Disorders – Fourth edition (DSM-IV) algorithms with the CIDI (Composite International Diagnostic Interview) and being aged between 18 and 70 years. | Severe internal or neurological disorders, MRI contraindications, use of antidepressants or other psychoactive medication (with the exception of stable benzodiazepine use for patients), lifetime diagnosis of psychotic disorders, bipolar disorder or personality disorders and dependence on drugs or alcohol. |

| | | | | |
|---|---|---|---|---|
| **MPIP** | Germany | M-CIDI/SCAN interview | M. A. R. S. sample: both first and recurrent episodes; RUD sample: only recurrent episodes with some patients scanned in remission | Munich Antidepressant Response Signature (MARS) study MDD subjects (clinical consensus diagnosis or M-CIDI (since 2008)): depressive syndromes secondary to any medical or neurological condition (e. g., intoxication, drug abuse, stroke), the presence of manic, hypomanic or mixed affective symptoms, lifetime diagnosis of alcohol dependence, illicit drug abuse or the presence of severe medical conditions (e.g., ischemic heart disease). Patients with bipolar depression were excluded for the current MR study. Control subjects: age > 65, MMSE<27, presence of severe somatic diseases or lifetime history of the following axis I disorders as assessed by the M-CIDI interview: alcohol dependence, drug abuse or dependence, possible psychotic disorder, mood disorder, anxiety disorder including OCD and PTSD, somatoform disorder, dissociative disorder NOS, and eating disorder 2. Recurrent unipolar depression (RUD) study: MDD subjects (SCAN interview): presence of manic episodes, mood incongruent psychotic symptoms, the presence of a lifetime diagnosis of intravenous drug abuse and depressive symptoms only secondary to alcohol or substance abuse or to medical illness or medication.Control subjects: presence of severe somatic diseases or life-time history of anxiety and affective disorders according to the Composite International Diagnostic-Screener (CIDI-S). All subjects: gross incidental MR findings such as territorial infarction, tumor, hydrocephalus, malformations and anatomical deviations (e.g. enlarged ventricles) that prevent appropriate image processing were additional exclusion criteria. RUD control samples were included. |

| | | | | |
|---|---|---|---|---|
| **Münster Neuroimaging Cohort** | Germany | SCID interview | Participants recruited by means of public advertisement and from the inpatient services. Inclusion criteria: age 16-65 years; patients were diagnosed with major depressive disorder by SCID-Interview | MDD subjects: presence of bipolar disorder, schizoaffective disorders and schizophrenia; substanc-erelated disorders or current benzodiazepine treatment (wash out of at least three half-lives before study participation), and former electroconvulsive therapy. Control subjects: any current or former psychiatric disorder. Both groups: any neurological abnormalities, MRI contra-indications |
| **NESDA** | Netherlands | CIDI interview | DSM-4 based diagnosis of MDD (6 month recency), using CIDI interview. 93 (60%) MDD patients have a comorbid ANX diagnosis. Age range 18-65 | |
| **QTIM** | Australia | CIDI interview | Retrospective questionnaire about depression episodes combined with an MRI study. The best described MDD episode is defined as the worst one (according to individuals). We have up to 5 supplementary episodes (briefly) described. Sample composed of twins and relatives. Population-based sample | MDD subjects: presence of axis-I disorders other than MDD and anxiety disorders Control subjects: antidepressant use, psychiatric disorders All subjects: relatedness between subjects, left handedness, history of neurological or other severe medical illness, head injury or current or past diagnosis of substance abuse, use of cognition affecting medication and general MRI contraindications |
| **SHIP-START** | Germany | M-CIDI interview | Population based longitudinal cohort study | MDD subjects: presence of axis-I disorders other than MDD, anxiety disorders, conversion, somatization and eating disorder. Control subjects: no lifetime diagnosis of depression, no antidepressiva, and severity index=0 All subjects: We removed subjects with medical conditions (e.g. a history of cerebral tumor, stroke, Parkinson’s diseases, multiple sclerosis, epilepsy, hydrocephalus, enlarged ventricles, pathological lesions) or due to technical reasons (e.g. severe movement artifacts or inhomogeneity of the magnetic |

| | | | | |
|---|---|---|---|---|
| | | | | field). |
| **SHIP-TREND** | Germany | M-CIDI interview | Population based longitudinal cohort study | MDD subjects: no special exclusion criteria Control subjects: no lifetime diagnosis of depression, no antidepressiva, and severity index=0 All subjects: We removed subjects with due to medical conditions (e.g. a history of cerebral tumor, stroke, Parkinson's diseases, multiple sclerosis, epilepsy, hydrocephalus, enlarged ventricles, pathological lesions) or due to technical reasons (e.g. severe movement artifacts or inhomogeneity of the magnetic field). |
| **Sydney** | Australia | SCID interview | | MDD subjects: presence of axis-I disorders other than MDD, panic disorder, social anxiety disorder, or generalized anxiety disorder. Control subjects: no Axis-I diagnosis, no medication use. Exclusion criteria for all subjects included medical instability (as determined by a psychiatrist), history of neurological disease (e.g. tumour, head trauma, epilepsy), medical illness known to impact cognitive and brain function (e.g. cancer), intellectual and/or developmental disability and insufficient English for neuropsychological assessment. All subjects were asked to abstain from drug or alcohol use for 48 hours prior to testing and informed about a drug screen protocol. |

**Supplementary Table S2.** Information on image acquisition parameters, software descriptions, and quality control.

| Cohort | Country | Scanner type | Sequence T1 | FreeSurfer version | Slice orientation | Operating system |
|---|---|---|---|---|---|---|
| **BiDirect** | Germany | 3 T Philips Intera scanner | 3D T1-weighted turbo field echo images were collected with a the following parameters: TR = 7.26, TE = 3.56, 9° flip angle, 160 sagittal slices, matrix dimension 256 x 256, FOV = 256 x 256mm, 2mm slice thickness (reconstructed to 1mm) and a resulting voxel size of 1x1x1mm | 5.3 | Sagittal | |
| **CliNG** | Germany | 3T Siemens Tim Trio | T1-weighted 3D MPRAGE; TR/TE/TI/FA=2250 ms/3.26 ms/900 ms/9°; image matrix = 256 x 256; 192 sagittal slices; voxel size= 1 mm3 | 5.3 | Sagittal | Linux |
| **CSAN** | Sweden | 3T Siemens MAGNETOM PRISMA | Whole-head t1-weighted MPRAGE (TR = 2300 ms, TE = 2.34 ms, FOV 250 × 250 mm, voxel size = 0.9 × 0.868 × 0.868 mm, flip angle = 8°) | 7.2 | Sagittal | Ubuntu |
| **FOR2107 - Marburg** | Germany | 3T Siemens Magnetom TiroTim syngo MR B17 | Sequence: 3D T1-weighted magnetization prepared rapid acquisition gradient echo (MPRAGE) - Sagittal Acquisition Direction, # of Slices 176, 0.5mm Slice Gap, 1.0x1.0x1.0 Voxel Size (mm3), TI 900 ms, TE 2.26 ms, TR 1900 ms, Flip Angle 9. | 5.3 | Sagittal | Red Hat Enterprise Linux Server release 5.11 (Tikanga) |
| **FOR2017 - Münster** | Germany | 3T Siemens PRISMA | Sequence: 3D T1-weighted magnetization prepared rapid acquisition gradient echo (MPRAGE). - Sagittal Acquisition Direction, # of Slices 192, 0mm Slice Gap, 1.0x1.0x1.0 Voxel Size (mm3), TI 900 ms, TE 2.28 ms, TR 1900 ms, Flip Angle 8 | 5.3 | Sagittal | Red Hat Enterprise Linux Server release 5.11 (Tikanga) |

| | | | | | | |
|---|---|---|---|---|---|---|
| **MOTAR** | Netherlands | 3T Philips Achieva | 32-channel head coil at the Spinoza centre. 3D Turbo Field Echo (TFE) T1-weighted structural MRI with scan parameters: TR = 8.1 ms, TE = 3.7 ms, flip angle = 8 degrees, matrix size = 240 x 240, 1 mm3 isotropic voxels. | 6.0 | Transverse (Axial) | Linux |
| **MPIP** | Germany | 1.5T GE and Siemens (the latter: only few cases) | #1: T1-weighted SPGR sagittal 3D volume. TR=1030 msec; TE=3.4 msec; 124 slices; matrix=256x256; FOV=23.0x23.0 cm2; voxel size=0.8975 mm x0.8975 mm x 1.2- 1.4 mm; flip angle=90°; birdcage resonator. #2: same scanner as #1, platform update Signa Excite, sagittal T1-weighted (spin echo sequence, TR=9.7 msec, TE=2.1 msec; FOV=25.0x25.0 cm2, voxel size=0.875 mm x0.875 mm x1.2 mm, 124- 132 slices, flip angle=90°. #3: Siemens 1.5 Tesla, Vario, 3D MPRAGE, TR=11.6 msec; TE=4.9 msec; FOV 23x23 cm2; matrix 512x512; 126 axial slices; voxel site 0.45 mm x 0.45 mm x 1.5 mm. (only N=2 subjects) | 5.3 | 1.5 GE: sagittal. 1.5 Siemens: axial | Linux 2.6.37.1-1.2- desktop x86_64 |
| **Münster Neuroimaging Cohort** | Germany | 3T Philips Gyroscan Intera | 3D fast gradient echo sequence (turbo field echo), repetition time = 7.4 milliseconds, echo time = 3.4 milliseconds, flip angle = 9°, two signal averages, inversion prepulse every 814.5 milliseconds, acquired over a field of view of 256 (feet -head [FH]) × 204 (anterior -posterior [AP]) × 160 (right -left [RL]) mm, phase encoding in AP and RL direction, reconstructed to cubic voxels of .5 mm × | 5.3 | Sagittal | Red Hat Enterprise Linux Server release 5.11 (Tikanga) |

| | | | | | | |
|---|---|---|---|---|---|---|
| | | | .5 mm × .5 mm | | | |
| **NESDA** | Netherlands | 3T Philips Achieva/Intera | 3D gradient-echo T1-weighted sequence. TR=9 msec; TE=3.5 msec; flip angle 8º, FOV = 256 mm; matrix: 25x62x56; in plane voxel size = 1 mm × 1 mm x 1 mm; 170 slices. | 5.0 | Sagittal | Linux |
| **QTIM** | Australia | Bruker 4T Wholebody MRI | 3D T1 weighted sequence. TR=1500 msec; TE=3.35 msec; flip angle=8°, 256 or 240 (coronal or sagittal) slices, FOV=240 mm, matrix 256x256x256 (or 256x256x240) | 5.1 | Coronal, then sagittal following software upgrade. | Linux- centos4_x86_64-stable-pub-v5.1.0 |
| **SHIP** | Germany | 1.5T Siemens Avanto | 3D T1-weighted (MP-RAGE/ axial plane); TR=1900 msec; TE=3.4 msec; Flip angle=15°; voxel size 1 mm x 1 mm x 1 mm | 5.3 (cortical), 5.1 (subcortical) | Axial | Centos6_x86_64 |
| **SHIP/TREND** | Germany | 1.5T Siemens Avanto | 3D T1-weighted (MP-RAGE/ axial plane); TR=1900 msec; TE=3.4 msec; Flip angle=15°; voxel size 1 mm x 1 mm x 1 mm | 5.3 (cortical), 5.1 (subcortical) | Axial | Centos6_x86_64 |
| **Sydney** | Australia | 3T GE MR750 | 3D T1-weighted sequence. TR=7.2 msec; TE=2.78 msec; matrix =256; FOV=240; No. slices=196; thick=0.9mm; inplane resolution=0.9375 | 5.1 | Coronal | Linux_Ubuntu16.04 lts 64bit |

**Supplementary Table S3. Performance accuracy of the brain age prediction model.**

| Group | r | $R^2$ | MAE (SD) in years | $_w$MAE | RMSE | Brain age gap (SD) in years |
|---|---|---|---|---|---|---|
| **Control** | 0.84 | 0.71 | 6.59 (5.01) | 0.11 | 8.28 | -0.04 (8.28) |
| **MDD** | 0.75 | 0.53 | 7.18 (5.50) | 0.12 | 9.05 | 1.33 (8.95) |

Abbreviations: r, Pearson's correlation coefficient; $R^2$, explained variance; MAE, Mean Absolute Error; $_w$MAE, weighted MAE (MAE/age range); RMSE, root mean squared error.

**Supplementary Table S4. Validation of PRS for MDD**

| Threshold | β | SE | OR | Upper CI | Lower CI | z-value | P-value | Significance |
|---|---|---|---|---|---|---|---|---|
| **0.00000005** | 0.042 | 0.035 | 1.043 | -0.027 | 0.110 | 1.200 | 0.230 | ns |
| **0.000001** | 0.047 | 0.035 | 1.048 | -0.021 | 0.116 | 1.353 | 0.176 | ns |
| **0.0002** | 0.072 | 0.035 | 1.075 | 0.004 | 0.141 | 2.067 | **0.039** | * |
| **0.001** | 0.11 | 0.036 | 1.116 | 0.040 | 0.180 | 3.070 | **0.002** | ** |
| **0.01** | 0.151 | 0.038 | 1.163 | 0.076 | 0.226 | 3.938 | **8.2×10$^{-6}$** | **** |
| **0.05** | 0.158 | 0.042 | 1.171 | 0.077 | 0.240 | 3.800 | **0.0001** | **** |
| **0.1** | **0.159** | **0.040** | **1.173** | **0.081** | **0.237** | **3.997** | **6.4×10$^{-5}$** | **** |
| **0.2** | 0.146 | 0.040 | 1.157 | 0.068 | 0.223 | 3.668 | **0.0002** | **** |
| **0.5** | 0.135 | 0.040 | 1.145 | 0.057 | 0.214 | 3.363 | **0.001** | *** |
| **1** | 0.141 | 0.041 | 1.151 | 0.061 | 0.221 | 3.442 | **0.001** | *** |

Models were residualized for 10 genetic PCs, age, sex, and scanning site.

**Supplementary Table S5. Validation of PRS for BMI**

| Threshold | β | SE | lower CI | upper CI | DF | t-value | P-value | Significance |
|---|---|---|---|---|---|---|---|---|
| **0.00000005** | 0.116 | 0.018 | 0.081 | 0.151 | 2701 | 6.444 | **1.4×10$^{-10}$** | **** |
| **0.000001** | 0.144 | 0.018 | 0.108 | 0.179 | 2701 | 8.003 | **1.8×10$^{-15}$** | **** |
| **0.0002** | 0.167 | 0.018 | 0.132 | 0.202 | 2701 | 9.374 | **1.4×10$^{-20}$** | **** |
| **0.001** | **0.189** | **0.018** | **0.154** | **0.223** | **2701** | **10.616** | **8.0×10$^{-26}$** | **** |
| **0.01** | 0.136 | 0.018 | 0.101 | 0.171 | 2701 | 7.583 | **4.6×10$^{-14}$** | **** |
| **0.05** | 0.101 | 0.018 | 0.066 | 0.137 | 2701 | 5.628 | **2.0×10$^{-8}$** | **** |
| **0.1** | 0.112 | 0.018 | 0.077 | 0.148 | 2701 | 6.238 | **5.1×10$^{-10}$** | **** |
| **0.2** | 0.111 | 0.018 | 0.076 | 0.146 | 2701 | 6.167 | **8.0×10$^{-10}$** | **** |
| **0.5** | 0.069 | 0.018 | 0.033 | 0.104 | 2701 | 3.802 | **0.0001** | **** |
| **1** | 0.067 | 0.018 | 0.031 | 0.102 | 2701 | 3.689 | **0.0002** | **** |

Models were residualized for 10 genetic PCs, age, sex, and scanning site.

**Supplementary Table S6. Associations between the brain age gap and PRS for MDD**

| Threshold | β | SE | lower CI | upper CI | DF | t-value | P-value | Significance |
|---|---|---|---|---|---|---|---|---|
| **0.00000005** | 0.023 | 0.013 | -0.003 | 0.048 | 3913 | 1.765 | 0.078 | ns |
| **0.000001** | 0.024 | 0.013 | -0.001 | 0.049 | 3913 | 1.865 | 0.062 | ns |
| **0.0002** | 0.029 | 0.013 | 0.004 | 0.054 | 3913 | 2.259 | **0.024** | * |
| **0.001** | 0.038 | 0.013 | 0.013 | 0.063 | 3913 | 2.939 | **0.003** | ** |
| **0.01** | **0.038** | **0.013** | **0.013** | **0.064** | **3913** | **2.979** | **0.003** | ** |
| **0.05** | 0.038 | 0.013 | 0.012 | 0.063 | 3913 | 2.927 | **0.003** | ** |
| **0.1** | 0.029 | 0.013 | 0.004 | 0.054 | 3913 | 2.234 | **0.026** | * |
| **0.2** | 0.029 | 0.013 | 0.004 | 0.055 | 3913 | 2.279 | **0.023** | * |
| **0.5** | 0.029 | 0.013 | 0.004 | 0.054 | 3913 | 2.247 | **0.025** | * |
| **1** | 0.031 | 0.013 | 0.006 | 0.056 | 3913 | 2.406 | **0.016** | * |

Models were residualized for 10 genetic PCs, age, age$^2$, sex, and scanning site.

**Supplementary Table S7. Associations between the brain age gap and PRS for BMI**

| Threshold | β | SE | lower CI | upper CI | DF | t-value | P-value | Significance |
|---|---|---|---|---|---|---|---|---|
| **0.00000005** | 0.002 | 0.013 | -0.023 | 0.027 | 3913 | 0.153 | 0.878 | ns |
| **0.000001** | 0.003 | 0.013 | -0.023 | 0.028 | 3913 | 0.213 | 0.831 | ns |
| **0.0002** | 0.013 | 0.013 | -0.013 | 0.038 | 3913 | 0.982 | 0.326 | ns |
| **0.001** | **0.029** | **0.013** | **0.003** | **0.054** | **3913** | **2.222** | **0.026** | * |
| **0.01** | 0.004 | 0.013 | -0.021 | 0.029 | 3913 | 0.299 | 0.765 | ns |
| **0.05** | -0.008 | 0.013 | -0.033 | 0.017 | 3913 | -0.632 | 0.527 | ns |
| **0.1** | -0.011 | 0.013 | -0.036 | 0.014 | 3913 | -0.839 | 0.401 | ns |
| **0.2** | -0.009 | 0.013 | -0.034 | 0.016 | 3913 | -0.702 | 0.483 | ns |
| **0.5** | -0.018 | 0.013 | -0.043 | 0.007 | 3913 | -1.394 | 0.163 | ns |
| **1** | -0.018 | 0.013 | -0.043 | 0.007 | 3913 | -1.411 | 0.158 | ns |

Models were residualized for 10 genetic PCs, age, age$^2$, sex, and scanning site.

**Supplementary Table S8. Associations between the brain age gap and PRS for CRP**

| Threshold | β | SE | lower CI | upper CI | DF | t-value | P-value | Significance |
|---|---|---|---|---|---|---|---|---|
| **0.00000005** | 0.010 | 0.013 | -0.015 | 0.035 | 3913 | 0.790 | 0.430 | ns |
| **0.000001** | 0.011 | 0.013 | -0.015 | 0.036 | 3913 | 0.817 | 0.414 | ns |
| **0.0002** | 0.011 | 0.013 | -0.014 | 0.037 | 3913 | 0.878 | 0.380 | ns |
| **0.001** | 0.022 | 0.013 | -0.004 | 0.047 | 3913 | 1.686 | 0.092 | ns |
| **0.01** | 0.032 | 0.013 | 0.007 | 0.057 | 3913 | 2.489 | **0.013** | * |
| **0.05** | 0.044 | 0.013 | 0.019 | 0.069 | 3913 | 3.400 | **0.001** | *** |
| **0.1** | 0.045 | 0.013 | 0.019 | 0.07 | 3913 | 3.457 | **0.001** | *** |
| **0.2** | 0.042 | 0.013 | 0.017 | 0.068 | 3913 | 3.276 | **0.001** | *** |
| **0.5** | 0.035 | 0.013 | 0.010 | 0.060 | 3913 | 2.730 | **0.006** | ** |
| **1** | 0.031 | 0.013 | 0.006 | 0.057 | 3913 | 2.438 | **0.015** | * |

Models were residualized for 10 genetic PCs, age, age$^2$, sex, and scanning site.

**Supplementary Table S9. Predictor-by-diagnosis interaction on the brain age gap.**

| Predictor | β | SE | DF | t-value | P-value | $P_{FDR}$ |
|---|---|---|---|---|---|---|
| **PRS MDD** | 0.009 | 0.026 | 3911 | 0.365 | 0.715 | 0.868 |
| **PRS BMI** | 0.025 | 0.026 | 3911 | 0.957 | 0.339 | 0.593 |
| **PRS CRP** | 0.008 | 0.026 | 3911 | 0.327 | 0.744 | 0.868 |
| **BMI** | -0.001 | 0.032 | 2698 | -0.042 | 0.966 | 0.966 |
| **Childhood trauma (CTQ total)** | 0.051 | 0.04 | 2194 | 1.285 | 0.199 | 0.593 |
| **Smoking (Current Status)** | 0.095 | 0.074 | 2589 | 1.281 | 0.200 | 0.593 |
| **Education (Years)** | -0.041 | 0.039 | 1916 | -1.061 | 0.289 | 0.593 |

PRS scores were residualized for 10 genetic PCs. Age, age$^2$, and sex were added as covariates in the model. Random intercepts for scanning sites were included.

**Supplementary Table S10. Validation of PRS for GrimAge acceleration residuals**

| Threshold | β | SE | lower CI | upper CI | DF | t-value | P-value | Significance |
|---|---|---|---|---|---|---|---|---|
| **0.00000005** | **0.075** | **0.035** | **0.006** | **0.143** | **806** | **2.126** | **0.034** | ***** |
| **0.000001** | 0.032 | 0.035 | -0.037 | 0.101 | 806 | 0.912 | 0.362 | ns |
| **0.0002** | 0.061 | 0.035 | -0.007 | 0.130 | 806 | 1.751 | 0.080 | ns |
| **0.001** | 0.064 | 0.035 | -0.004 | 0.133 | 806 | 1.834 | 0.067 | ns |
| **0.01** | 0.094 | 0.035 | 0.025 | 0.163 | 806 | 2.685 | **0.007** | ** |
| **0.05** | 0.097 | 0.035 | 0.028 | 0.165 | 806 | 2.758 | **0.006** | ** |
| **0.1** | 0.108 | 0.035 | 0.039 | 0.176 | 806 | 3.086 | **0.002** | ** |
| **0.2** | 0.076 | 0.035 | 0.007 | 0.145 | 806 | 2.174 | **0.030** | * |
| **0.5** | 0.065 | 0.035 | -0.004 | 0.133 | 806 | 1.838 | 0.066 | ns |
| **1** | 0.055 | 0.035 | -0.014 | 0.124 | 806 | 1.564 | 0.118 | ns |

Models were residualized for 10 genetic PCs, age, sex, and batch.

**Supplementary Table S11. Validation of PRS for Hannum age acceleration residuals**

| Threshold | β | SE | lower CI | upper CI | DF | t-value | P-value | Significance |
|---|---|---|---|---|---|---|---|---|
| **0.00000005** | 0.018 | 0.035 | -0.051 | 0.087 | 806 | 0.510 | 0.610 | ns |
| **0.000001** | 0.051 | 0.035 | -0.018 | 0.120 | 806 | 1.452 | 0.147 | ns |
| **0.0002** | **0.072** | **0.035** | **0.003** | **0.141** | **806** | **2.060** | **0.040** | ***** |
| **0.001** | 0.049 | 0.035 | -0.020 | 0.118 | 806 | 1.391 | 0.164 | ns |
| **0.01** | 0.058 | 0.035 | -0.010 | 0.127 | 806 | 1.663 | 0.097 | ns |
| **0.05** | 0.036 | 0.035 | -0.033 | 0.105 | 806 | 1.028 | 0.304 | ns |
| **0.1** | -0.023 | 0.035 | -0.092 | 0.046 | 806 | -0.643 | 0.521 | ns |
| **0.2** | -0.025 | 0.035 | -0.094 | 0.044 | 806 | -0.699 | 0.484 | ns |
| **0.5** | -0.019 | 0.035 | -0.088 | 0.050 | 806 | -0.546 | 0.585 | ns |
| **1** | -0.011 | 0.035 | -0.080 | 0.058 | 806 | -0.323 | 0.747 | ns |

Models were residualized for 10 genetic PCs, age, sex, and batch.

### Supplementary Table S12. Validation of PRS for Horvath age acceleration residuals

| Threshold | β | SE | lower CI | upper CI | DF | t-value | P-value | Significance |
|---|---|---|---|---|---|---|---|---|
| **0.00000005** | 0.126 | 0.035 | 0.058 | 0.195 | 806 | 3.617 | **0.000** | *** |
| **0.000001** | 0.159 | 0.035 | 0.091 | 0.227 | 806 | 4.586 | **0.000** | **** |
| **0.0002** | **0.169** | **0.035** | **0.101** | **0.236** | **806** | **4.859** | **0.000** | **** |
| **0.001** | 0.146 | 0.035 | 0.078 | 0.214 | 806 | 4.194 | **0.000** | **** |
| **0.01** | 0.135 | 0.035 | 0.066 | 0.203 | 806 | 3.864 | **0.000** | *** |
| **0.05** | 0.091 | 0.035 | 0.022 | 0.159 | 806 | 2.587 | **0.010** | ** |
| **0.1** | 0.082 | 0.035 | 0.013 | 0.151 | 806 | 2.338 | **0.020** | * |
| **0.2** | 0.085 | 0.035 | 0.016 | 0.154 | 806 | 2.426 | **0.015** | * |
| **0.5** | 0.077 | 0.035 | 0.008 | 0.145 | 806 | 2.183 | **0.029** | * |
| **1** | 0.081 | 0.035 | 0.012 | 0.149 | 806 | 2.300 | **0.022** | * |

Models were residualized for 10 genetic PCs, age, sex, and batch.

### Supplementary Table S13. Validation of PRS for PhenoAge age acceleration residuals

| Threshold | β | SE | lower CI | upper CI | DF | t-value | P-value | Significance |
|---|---|---|---|---|---|---|---|---|
| **0.00000005** | **0.160** | **0.035** | **0.092** | **0.228** | **806** | **4.604** | **0.000** | **** |
| **0.000001** | 0.151 | 0.035 | 0.083 | 0.219 | 806 | 4.343 | **0.000** | **** |
| **0.0002** | 0.123 | 0.035 | 0.054 | 0.191 | 806 | 3.519 | **0.000** | *** |
| **0.001** | 0.078 | 0.035 | 0.009 | 0.146 | 806 | 2.214 | **0.027** | * |
| **0.01** | 0.072 | 0.035 | 0.003 | 0.141 | 806 | 2.052 | **0.040** | * |
| **0.05** | 0.084 | 0.035 | 0.015 | 0.153 | 806 | 2.397 | **0.017** | * |
| **0.1** | 0.056 | 0.035 | -0.012 | 0.125 | 806 | 1.605 | 0.109 | ns |
| **0.2** | 0.058 | 0.035 | -0.011 | 0.127 | 806 | 1.657 | 0.098 | ns |
| **0.5** | 0.073 | 0.035 | 0.004 | 0.141 | 806 | 2.071 | **0.039** | * |
| **1** | 0.069 | 0.035 | 0.000 | 0.138 | 806 | 1.966 | **0.050** | * |

Models were residualized for 10 genetic PCs, age, sex, and batch.

### Supplementary Table S14. Associations between the brain age gap and epigenetic clocks

| Epigenetic aging | β | SE | lower CI | upper CI | DF | t-value | P-value | $P_{FDR}$ |
|---|---|---|---|---|---|---|---|---|
| **GrimAge acceleration residuals** | 0.139 | 0.031 | 0.079 | 0.199 | 801 | 4.527 | **$6.9\times10^{-6}$** | **$2.8\times10^{-5}$** |
| **Hannum age acceleration residuals** | 0.043 | 0.031 | -0.017 | 0.102 | 801 | 1.393 | 0.164 | 0.164 |
| **Horvath age acceleration residuals** | 0.053 | 0.030 | -0.006 | 0.111 | 801 | 1.767 | 0.078 | 0.104 |
| **PhenoAge acceleration residuals** | 0.070 | 0.030 | 0.012 | 0.128 | 801 | 2.359 | **0.019** | **0.037** |

Models were residualized for age, age$^2$, sex, and batch.

**Supplementary Table S15. Associations between the brain age gap and PRS for GrimAge acceleration residuals**

| Threshold | β | lower CI | upper CI | SE | DF | t-value | P-value | Significance |
|---|---|---|---|---|---|---|---|---|
| **0.00000005** | 0.007 | -0.018 | 0.033 | 0.013 | 3913 | 0.578 | 0.563 | ns |
| **0.000001** | 0.007 | -0.018 | 0.032 | 0.013 | 3913 | 0.544 | 0.587 | ns |
| **0.0002** | 0.001 | -0.024 | 0.026 | 0.013 | 3913 | 0.09 | 0.928 | ns |
| **0.001** | -0.012 | -0.037 | 0.014 | 0.013 | 3913 | -0.9 | 0.368 | ns |
| **0.01** | 0.000 | -0.026 | 0.025 | 0.013 | 3913 | -0.025 | 0.980 | ns |
| **0.05** | 0.008 | -0.017 | 0.033 | 0.013 | 3913 | 0.624 | 0.532 | ns |
| **0.1** | 0.013 | -0.012 | 0.039 | 0.013 | 3913 | 1.037 | 0.300 | ns |
| **0.2** | 0.013 | -0.013 | 0.038 | 0.013 | 3913 | 0.974 | 0.330 | ns |
| **0.5** | 0.010 | -0.015 | 0.035 | 0.013 | 3913 | 0.777 | 0.437 | ns |
| **1** | 0.014 | -0.012 | 0.039 | 0.013 | 3913 | 1.055 | 0.291 | ns |

Models were residualized for 10 genetic PCs, age, age$^2$, sex, and scanning site.

**Supplementary Table S16. Associations between the brain age gap and PRS for Hannum age acceleration residuals**

| Threshold | β | lower CI | upper CI | SE | DF | t-value | P-value | Significance |
|---|---|---|---|---|---|---|---|---|
| **0.00000005** | -0.012 | -0.037 | 0.013 | 0.013 | 3913 | -0.934 | 0.350 | ns |
| **0.000001** | -0.023 | -0.048 | 0.003 | 0.013 | 3913 | -1.762 | 0.078 | ns |
| **0.0002** | -0.018 | -0.044 | 0.007 | 0.013 | 3913 | -1.423 | 0.155 | ns |
| **0.001** | -0.013 | -0.039 | 0.012 | 0.013 | 3913 | -1.031 | 0.303 | ns |
| **0.01** | 0.024 | -0.001 | 0.049 | 0.013 | 3913 | 1.856 | 0.064 | ns |
| **0.05** | 0.016 | -0.010 | 0.041 | 0.013 | 3913 | 1.208 | 0.227 | ns |
| **0.1** | 0.014 | -0.011 | 0.039 | 0.013 | 3913 | 1.095 | 0.274 | ns |
| **0.2** | 0.023 | -0.002 | 0.049 | 0.013 | 3913 | 1.819 | 0.069 | ns |
| **0.5** | 0.028 | 0.002 | 0.053 | 0.013 | 3913 | 2.142 | **0.032** | * |
| **1** | **0.030** | **0.005** | **0.055** | **0.013** | **3913** | **2.332** | **0.020** | * |

Models were residualized for 10 genetic PCs, age, age$^2$, sex, and scanning site.

**Supplementary Table S17. Associations between the brain age gap and PRS for Horvath age acceleration residuals**

| Threshold | β | lower CI | upper CI | SE | DF | t-value | P-value | Significance |
|---|---|---|---|---|---|---|---|---|
| **0.00000005** | **-0.037** | **-0.062** | **-0.012** | **0.013** | **3913** | **-2.884** | **0.004** | ** |
| **0.000001** | -0.034 | -0.059 | -0.009 | 0.013 | 3913 | -2.626 | **0.009** | ** |
| **0.0002** | -0.014 | -0.039 | 0.012 | 0.013 | 3913 | -1.059 | 0.290 | ns |
| **0.001** | -0.008 | -0.033 | 0.017 | 0.013 | 3913 | -0.608 | 0.543 | ns |
| **0.01** | 0.003 | -0.023 | 0.028 | 0.013 | 3913 | 0.201 | 0.841 | ns |
| **0.05** | 0.000 | -0.025 | 0.026 | 0.013 | 3913 | 0.017 | 0.986 | ns |
| **0.1** | 0.005 | -0.021 | 0.030 | 0.013 | 3913 | 0.367 | 0.714 | ns |
| **0.2** | -0.001 | -0.027 | 0.024 | 0.013 | 3913 | -0.106 | 0.916 | ns |
| **0.5** | 0.000 | -0.025 | 0.025 | 0.013 | 3913 | -0.008 | 0.994 | ns |
| **1** | 0.000 | -0.025 | 0.025 | 0.013 | 3913 | 0.010 | 0.992 | ns |

Models were residualized for 10 genetic PCs, age, age$^2$, sex, and scanning site.

**Supplementary Table S18. Associations between the brain age gap and PRS for PhenoAge acceleration residuals**

| Threshold | β | lower CI | upper CI | SE | DF | t-value | P-value | Significance |
|---|---|---|---|---|---|---|---|---|
| **0.00000005** | -0.001 | -0.026 | 0.025 | 0.013 | 3913 | -0.055 | 0.956 | ns |
| **0.000001** | 0.005 | -0.021 | 0.030 | 0.013 | 3913 | 0.367 | 0.713 | ns |
| **0.0002** | -0.007 | -0.032 | 0.018 | 0.013 | 3913 | -0.543 | 0.587 | ns |
| **0.001** | -0.003 | -0.028 | 0.022 | 0.013 | 3913 | -0.242 | 0.809 | ns |
| **0.01** | -0.010 | -0.035 | 0.016 | 0.013 | 3913 | -0.753 | 0.451 | ns |
| **0.05** | -0.001 | -0.027 | 0.024 | 0.013 | 3913 | -0.108 | 0.914 | ns |
| **0.1** | -0.002 | -0.027 | 0.023 | 0.013 | 3913 | -0.171 | 0.864 | ns |
| **0.2** | 0.000 | -0.026 | 0.025 | 0.013 | 3913 | -0.034 | 0.973 | ns |
| **0.5** | -0.006 | -0.031 | 0.020 | 0.013 | 3913 | -0.440 | 0.660 | ns |
| **1** | -0.002 | -0.027 | 0.023 | 0.013 | 3913 | -0.145 | 0.885 | ns |

Models were residualized for 10 genetic PCs, age, age$^2$, sex, and scanning site.
